# Sequencing and analysis of *Arabidopsis thaliana* NOR2 reveal its distinct organization and tissue-specific expression of rRNA ribosomal variants

**DOI:** 10.1101/2020.09.10.272005

**Authors:** Jason Sims, Giovanni Sestini, Christiane Elgert, Arndt von Haeseler, Peter Schlögelhofer

## Abstract

Despite vast differences between organisms, some characteristics of their genomes are conserved, such as the nucleolus organizing region (NOR). The NOR is constituted of multiple, highly repetitive rDNA genes, encoding the catalytic ribosomal core RNAs which are transcribed from 45S rDNA units. Their precise sequence information and organization remained uncharacterized.

We used a combination of long- and short-read sequencing technologies to assemble contigs of the *Arabidopsis* NOR2 rDNA domain providing a first map. We identified several expressed rRNA gene variants which are integrated into translating ribosomes in a tissue-specific manner. These findings support the concept of tissue specific ribosome subpopulations that differ in their rRNA composition and provide the higher order organization of NOR2.

## Introduction

Ribosomes are one of the oldest and most complex molecular machines in extant life. In all living cells, they are responsible for translating the genetic information residing on mRNAs into proteins. Ribosomes are large complex molecules, build from at least 80 proteins (estimate for *Arabidopsis*) and four types of rRNAs that constitute the catalytic core ^1^. In *Arabidopsis* the 18S, 5.8S and 25S rRNA genes are organized, in a head to tail manner, within large domains known as nucleolus organizing regions (NORs) ^2,3^ located at the subtelomeric domains of chromosomes 2 and 4. A single rDNA unit contains three rRNA genes (18S, 5.8S and 25S) with two internal and external transcribed sequences (ITS, ETS). Additionally, each unit contains a core promoter sequence preceded by one or multiple spacer promoters flanked by *Sal*I repeat boxes ^4^. Several studies have shown that the rDNA repeats are not completely identical, with variability within the intergenic regions and to some extent also in the coding sequences ^5–7^. It remains unknown whether these variations are present in the translating ribosome and have a functional impact. Four main variations (VAR) are located within the 3′ ETS and grouped on separate chromosomes ^6,8^. VAR1 and VAR3a, which are believed to be generally interspersed, are located on NOR2 where as VAR2 and VAR3b-c are located on NOR4^8,9^.

Although the ribosomal RNA transcripts account for approximately 50% of all transcribed RNA in a cell, only a fraction of the tandemly repeated rDNA genes is transcribed at a given time ^6,7^. The transcriptional regulation of individual rRNA genes and their organization within the NOR domains remains unknown, for any organism, due to their repetitive nature that hinders classical sequencing approaches ^7,10^. In fact, it has never been possible to assemble the complete NOR by directly sequencing the genome of *Arabidopsis* or that of any organism. An in-depth understanding of the regulation and organization of the rDNA repeats would require the analysis of the full sequence of the NORs.

In this study, we used a combination of long- and short-read sequencing to obtain the exact sequence information of 405 individual rDNA repeats and their arrangement. We sequenced 59 BACs containing ~7 rDNA repeats each and devised a sophisticated barcoding system to align each BAC and build long rDNA contigs. This allowed us to assess the higher-order organization of *Arabidopsis thaliana’*s NOR2 and build its first sequence draft. We identified several sequence variations in the rRNA genes and established that the corresponding rRNAs are expressed and integrated into mature and translating ribosomes in a tissue-specific manner. These findings uncover a complex transcriptional regulation of rDNA genes and support the concept of tissue-specific ribosome subpopulations^11^ that differ in their rRNA composition.

## Results

### Sequencing of the NOR2 rDNA

Our aim was to generate a complete assembly of the NOR2 of *Arabidopsis thaliana*. In order to successfully sequence and build NOR2, we relied on a bacterial artificial chromosome (BAC) based approach ^12^ (Fig. 1a). BACs, initially generated to obtain the *Arabidopsis* physical map ^13^, containing rDNA repeats were ordered from the ABRC stock center ^14^. They were classified according to the type of 3′ ETS variant (VAR) in order to discriminate their origin from either NOR2 or NOR4 ^8^ (Suppl. Fig 1a). BACs containing rRNA genes with VAR1 or VAR3 were further processed and sequenced to construct a draft of the NOR 2 domain. Supercoiled BACs from bacterial cultures were linearized with the restriction enzyme *Apa*I prior to sequencing and analyzed by Pulse Field Gel Electrophoresis (PFGE) (Suppl. Fig 1b).

**Figure 1:**
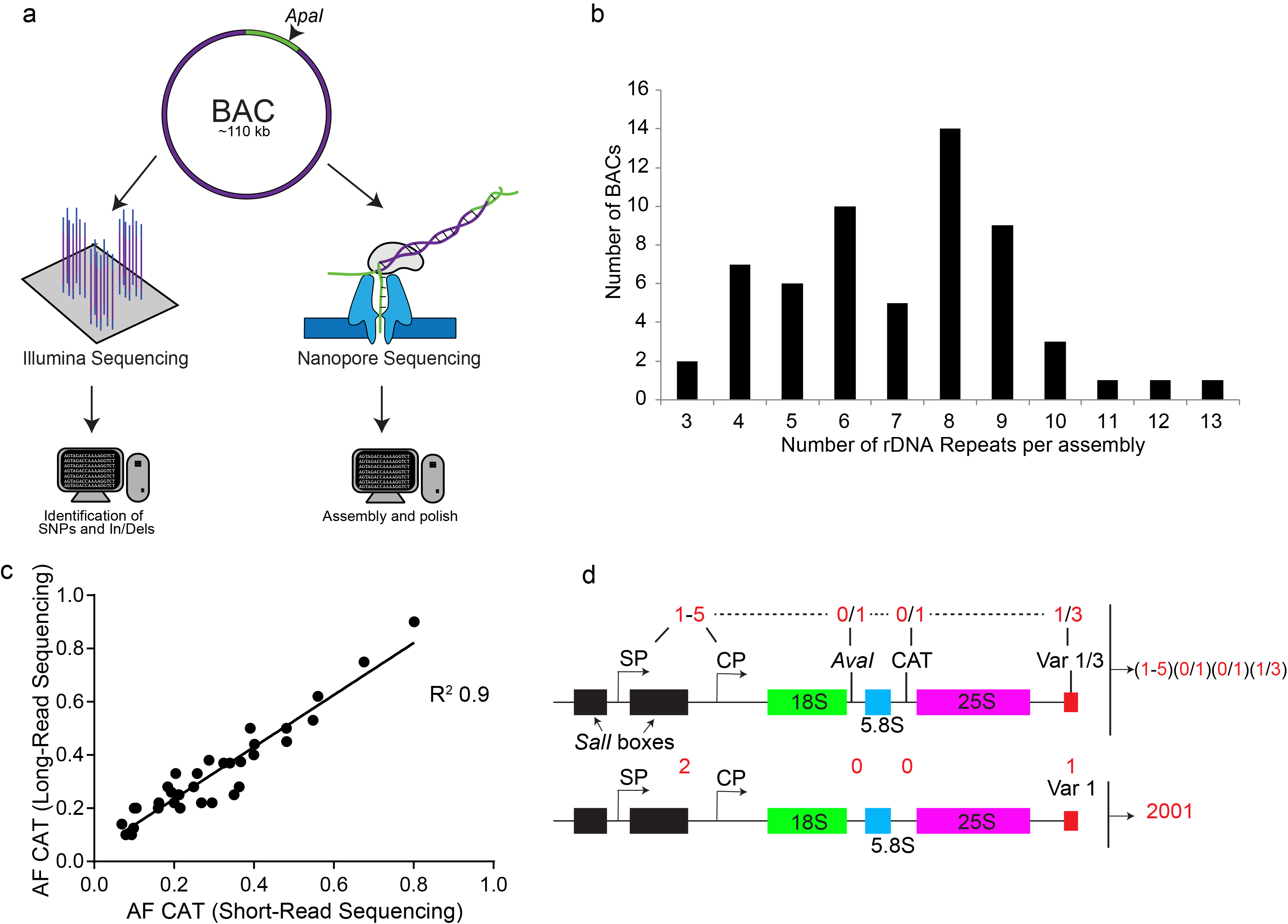
Sequencing and polishing BACs containing rDNA contigs. **a**, Scheme depicting the sequencing and assembly strategy combining long-read (NanoPore) and short-read sequencing (Illumina) of BACs containing rDNA genes. **b**, Graph depicting the number of rDNA units per assembled BAC. **c**, Correlation analysis of the CAT allele frequencies derived from the long- and short-read sequencing approaches. Each dot represents an individual assembly. **d**, Graphical description of the numeric codes attributed to the described rDNA unit features. Each four-digit code corresponds to a unique rDNA variant. An example is shown of an rDNA variant that carries <u>2</u> promoters (1 spacer promoter, SP and 1 core promoter, CP), no *Ava*I site (<u>0</u>), no CAT insertion (<u>0</u>) and the 3’ ETS VAR<u>1</u>. It is defined as a “<u>2001</u>” rDNA repeat.

In total we sequenced 59 BACs with an average BAC assembly size of about 77 kb (Table S1). We used reads longer than 50 kb for the assembly of the BACs (Suppl. Fig 1c). The assemblies contain an average of 7 rDNA repeats of ~10 kb each. The assembly sizes of 49 out of 59 BACs are coherent with the expected sizes according to the PFGE analysis ^14^ (Suppl. Fig. 1b) and contain the vector backbone separated at the *Apa*I cut site. These BACs were considered as fully assembled. The number of rDNA repeats per assembly is depicted in Figure 1b. In parallel, the 49 fully assembled BACs were also sequenced by short read sequencing technology (Illumina) in order to confirm single nucleotide polymorphisms (SNPs) and insertions and deletions (InDels) (Table S1). The Illumina reads of each BAC were mapped with the software bowtie2 ^15^ to a reference rDNA repeat (see Materials & Methods). SNPs and InDels were called with LoFreq ^16^, a base quality–aware algorithm designed for conservative detection of rare sequence variants that implements a strand bias test with Bonferroni correction ^16^ (Table S2). The correctness of the long-read assemblies was confirmed by: (i) restriction digest with *Fsp*I and *Xho*I which releases the repetitive *Sal*I boxes, a prominent length polymorphism marker in the rDNA ^5^ (Suppl. Fig 1d); (ii) cleaved amplified polymorphic sequence (CAPS) analysis of a frequent polymorphism at position 4133 (position relates to reference rDNA) which is cut by the *Ava*I restriction enzyme ^7^ (Suppl. Fig 1e) and (iii) by comparing the nature and the numbers of SNPs and InDels in a given assembly with the SNP/InDel allele frequencies (AFs) derived by short-read Illumina sequencing (Fig.1c, Suppl. Fig. 1f, Table S1). In the assemblies analyzed, the experimentally determined length of the *Sal*I boxes matched the expected sizes (Suppl. Fig. 1d). BAC assembly F2C3 after restriction digest released fragments of ~4700 and ~3900 bp, F1E12 and F2G13 released fragments of ~3500bp and ~1500 bp and F2G18 released fragments of ~1500 bp (Suppl. Fig. 1d). Furthermore, the CAPS analysis confirmed the presence of the polymorphism at the sites anticipated from the assemblies (Suppl. Fig. 1e). Finally, for the full assemblies (49 BACs) the AFs determined by short-read sequencing matched in number and nature those determined in the assembled BACs. For instance, the R^2^ correlation between the SNP/InDel frequencies of the long-read assemblies with the short-read sequences for the *Ava*I restriction site and the trinucleotide insertion “CAT” polymorphisms ^7^ (position 4466 of the reference rDNA) is 0.74 and 0.9, respectively (Fig.1c, Suppl. Fig. 1f, Table S1). In all assemblies the rDNA repeats are organized in a head-to-tail manner as shown previously ^3^. This latter finding substantiates the quality and reliability of our sequencing efforts, arguing against a possible re-arrangement of the multiple rDNA repeat units present in the BACs during library generation or faulty assembly. In order to confirm that the SNPs and InDels identified in the 49 fully assembled BACs were present in the corresponding genome of *A. thaliana* Col-0 plants, we made use of whole genome sequencing data previously generated in our lab (Kurzbauer et al. 2020 submitted). This data set was obtained by Illumina 125bp paired-end sequencing and yielded a genome-wide 30-fold coverage. The reads corresponding to the rDNA were mapped to the reference rDNA repeat (Table S3) and SNPs/InDels called with LoFreq. We obtained the nature and frequency of occurrence of SNPs and InDels of all rDNA units present in the genome, located on both NOR2 and NOR4. We only took into consideration allele frequencies above 0.125% (frequencies obtained if one SNP/InDel would be present in a single rDNA unit out of ~800 units) since it is estimated that the *Arabidopsis* genome contains ~800 rDNA repeats ^2^. 58% of the SNPs/InDels in the BACs were also present in the genome (Table S3, Table S4). The remaining 42% of SNPs/InDels that were not found in the genome could represent genetic background alterations which may have arisen from mutagenic events during the generation of the BAC library or are false positives called by LoFreq (Table S4). We were able to successfully assemble and confirm 49 BACs which contained an average of 7 rDNA units per BAC. These assemblies provided the base to build the first draft of the *Arabidopsis* NOR2.

### rRNA gene variants guide the assembly of NOR2

To guide the building of a first draft of the NOR 2 domain, we devised two “barcode” systems to discriminate each rDNA repeat unit, which are 98% identical. The first barcode system is based on four markers: (1) number of core and spacer promoters ^5^, (2) presence of the *Ava*I restriction site (at position 4133 of the reference rDNA) ^9^, (3) presence of a specific trinucleotide (CAT) insertion (at coordinate 4466 of the reference rDNA) ^9^, (4) length of the 3′ ETS ^6^. This barcode system allows us to individualize each rDNA unit and guide the overlap between BAC assemblies generating larger contigs. The second barcode system is based on the length and number of the *Sal*I boxes ^5^. It has been used separately due to the highly complex nature of the *Sal*I boxes.

The number of promoters, the length of the 3’ETS and the *Sal*I boxes were selected as markers since they largely vary between different rDNA repeats ^5,6,17^. The *Ava*I restriction site is equally distributed between NOR2 and NOR4 while the CAT marker was shown to be strongly enriched on NOR2 ^8^. All other SNPs/InDels where not taken into consideration since they are either of low allelic frequency or are associated to the markers mentioned above. The “four marker barcode system” shows that the most frequently occurring unit type is 2001 (38%) (2 promoters, absence of *Ava*I, absence of CAT and 3’ ETS variant 1) (Table 1), 11% are of type 2003, 10% are of type 3001, 8.6% are of type 3111 (Fig. 1d, Suppl. Fig 2a). For the “*Sal*I box barcode system” each rDNA repeat was categorized by an alphanumeric code with each letter corresponding to a specific *Sal*I box length. rDNA units with multiple letters indicate the presence of multiple *Sal*I boxes of a given type (Table 1). The most frequent *Sal*I box combinations are EZ (14%), EU (12.5%), EV (11%), EEV (8.5%), EEU (8.3%), FQ (6%), I (5.8%) and M (5.8%) (Suppl. Fig. 2b).

We analyzed the individual rDNA units and found that units containing the CAT insertion and the EEV or EEU *Sal*I box type are associated in 60% of all cases with the *AvaI* site. In contrast, rDNA units containing the CAT and the M or FQ *Sal*I box types are devoid of the *Ava*I site in all cases. Interestingly, the *Ava*I site is in 99% of all cases associated with the CAT insertion site (Suppl. Fig. 2c). A small fraction (≤5%) of the rDNA units contained large deletions spanning 270 to 334bp located within both the transcribed and non-transcribed regions (Suppl. Fig 3a-c.). These deletions are located within the 5′ ETS (A and B), the 18S rRNA ^18^ (C) and spanning the 18S and the ITS1 (D) (Suppl. Fig 2a, Suppl. Fig 3a-c). This initial sequence analysis and the establishment of the barcode systems gave each rDNA unit an identity and provided a solid base to overlap BAC assemblies with high confidence.

In order to determine possible overlapping BAC assemblies we first designed a program (OverBACer) that determines, for each assembly, the sizes of the *Sal*I boxes, the types of the 3’ETS variants present and the distances (in bps) between them. The program generates a string of marker identities and distances creating a simple and unique identifier for each assembly. Subsequently, the assembly-specific strings were used to find matches between the different assemblies. The matching overlaps were manually confirmed by evaluating the *Ava*I and CAT markers. OverBACer automatically generated 53 unique contigs using the sequencing input from 59 BACs, with an average length of ~80 kb. 10 assemblies were found to overlap with high confidence, with 230 kb as the longest contig (derived from BACs F1F17-F2C3-F2J17). The other contigs of 200, 90 and 80 kb derived from BACs F2N4-F1B23-F2G13, F2G18-F19A6 and F1F11-F1N7, respectively. The relative low amount of overlaps identified between the 59 BACs led to the assumption that the NOR2 is much larger than previously anticipated (~400 repeats) ^3^. The quality and length of the contigs generated provided a first map of NOR2 that could be analyzed for its higher-order organization albeit of being incomplete.

### NOR2 is organized in distinct rDNA unit clusters

Our sequencing results revealed that NOR2 is composed of heterogeneous rDNA units. We therefore evaluated whether the individual rDNA units, distinguished by their sequence features, are organized non-randomly. We designed a program (Neighbour Finder) that identifies rDNA units according to their features and counts the occurrence of neighboring rDNA unit types. The barcode system described above was used for the analysis: number of promoters (P, 1 to 5), presence (1) or absence (0) of the *Ava*I restriction site, presence (1) or absence (0) of the CAT polymorphism and type of the 3’ ETS variant (1 or 3) (Fig. 1d, Table 1). We classified rDNA units according to the selected features and analyzed co-occurring neighbors at 10, 20 or 30 kb distances, corresponding to the immediate neighboring unit, one further downstream and two further downstream, respectively (Fig. 3a, Suppl. File 1). A Monte Carlo simulation (see Material & Methods) of all rDNA units, maintaining the head-to-tail arrangement and the BAC configuration was generated as a control. Indeed, co-occurrences of some barcode pairs in the contigs are more frequent than expected from a random distribution. The analysis revealed that rDNA unit pairs (2011, 1011), (3111, 3001) and (2001, 2001) frequently co-occur in a distance of 10 kb and 20 kb. In contrast, rDNA unit pairs (2001, 3001) and (2001, 3111) never co-occur in a distance of 10 kb, 20 kb and 30 kb, while they do so in the random simulation (Fig 3b-e, Suppl. File 1).

These results underline that rDNA unit types are not randomly distributed along NOR2 and that they form clusters. To illustrate this, we plotted the frequency of co-occurring pairs identified at 10 kb distance against the frequency of corresponding pairs counted at 20 kb, 30 kb distance or in the random simulation. The R^2^ values clearly indicate correlations at 20 kb (R^2^ 0.59) and 30 kb (R^2^ 0.38), but not when compared to the frequency of pairs obtained from the Monte Carlo simulation (R^2^ 0.028) (Fig. 2f-h). Our results are in line with the findings obtained by Copenhaver et al. ^3^ that suggested the existence of rDNA sub-clusters.

**Figure 2:**
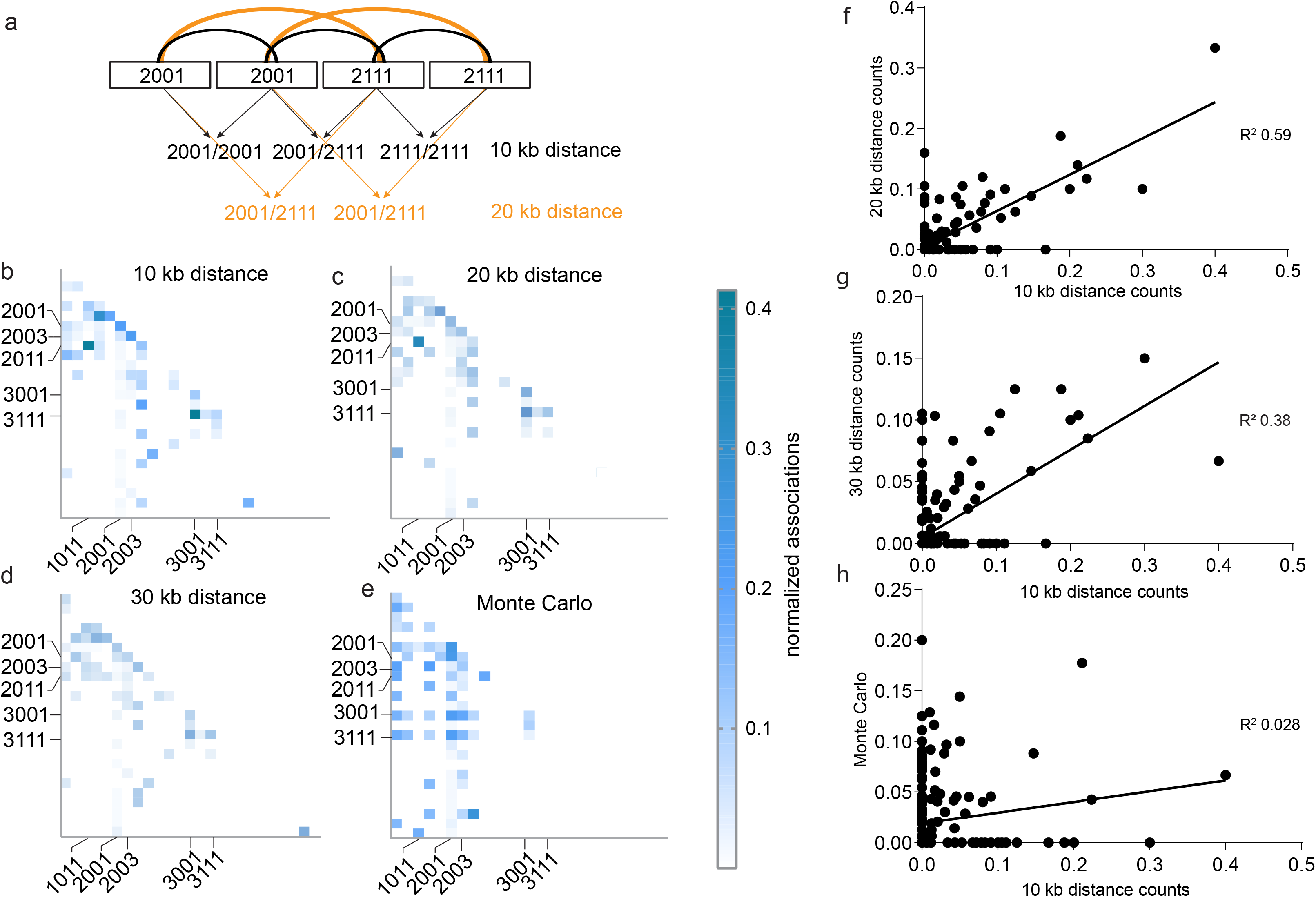
NOR2 is organized in distinct rDNA clusters. **a**, Graphical description of the NeighbourFinder program. It counts the occurrence of rDNA unit types at a 10kb (immediate neighbor), 20kb (one unit downstream) or 30kb (neighbor two units downstream) distance. **b-e**, Heat maps depicting the number of neighboring rDNA unit types at a 10 kb, 20 kb or 30 kb distance normalized to the sum of the rDNA units bearing the given features. Dark blue colors depict the highest relative number (0.42) and white the lowest (0). rDNA variants are listed as follows for each heat map: on the y-axis from top to bottom and for the x-axis from left to right: 1001, 1003, 1011, 1013, 1111, 2001, 2003, 2011, 2013, 2101, 2111, 2113, 3001, 3011, 3111, 4001, 4111, 5001, 1003D, 1011D, 1013D, 2001A, 2001C, 2001B, 2003C, (A, B, C, D depict large deletions, see Table S4 for glossary) Most abundant ones are indicated. **f-h**, Scatter plots showing the correlation between the frequencies of associated rDNA units of the 10kb distance with the 20kb distance (R^2^= 0.59) (**f**), 10kb distance with 30 kb distance (R^2^=0.38) (**g**), and the 10kb distance with the numbers obtained by the Monte Carlo simulation (R^2^=0.028) (**h**).

**Figure 3:**
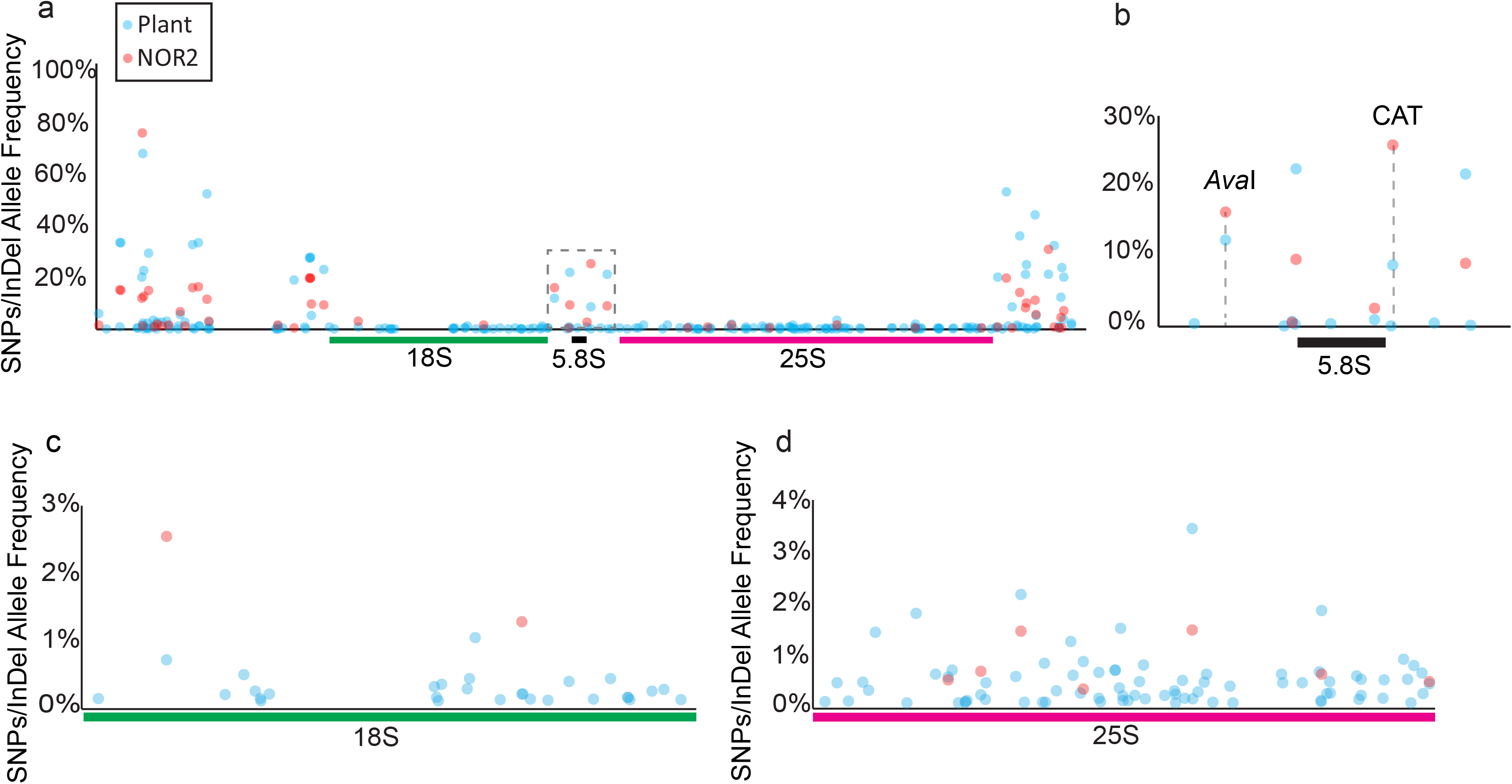
rDNA variants are differentially distributed between NOR2 and NOR4. **a**, Plot depicting the SNP/InDel allele frequencies in rDNA units derived from genomic plant DNA sequencing (blue dots; representing allele frequencies in both NOR2 and NOR4) and from BAC sequencing (red dots; representing allele frequencies only in NOR2). Dashed box indicates the enlarged area depicted in **b**. **b**, 5.8S rDNA region with the *Ava*I and the CAT alleles highlighted. **c**, Allele frequencies in the 18S rDNA region. **d** Allele frequencies in the 25S rDNA region.

We also performed such an analysis after classifying rDNA units according to their *Sal*I boxes. We find rDNA units harboring one, two or three *Sal*I boxes of a similar type in close proximity to one another at a 10, 20 (R^2^ 0.52) and 30 (R^2^ 0.38) kb distance (Suppl. Fig 4a). These results are different from the simulated data set where the units form distinct clusters not present in the experimental data (Suppl. Fig 4b-d). Accordingly, the frequencies of co-occurring *Sal*I boxes from the simulated data do not correlate with those obtained within a 10 kb distance (R^2^ 0.005) (Suppl. Fig 4d).

The rDNA units found to co-occur following the analysis with four markers correlate well with those identified by the analysis using the *Sal*I boxes. This represents a good internal control of our analysis as the number of promoters is proportional to the number of *Sal*I boxes, as also shown previously by Havlova et al.^5^ These results suggest a higher-order organization of NOR2 highlighting that rDNA units with similar features tend to form clusters.

The higher order organization and the analyses of the 405 rDNA units of NOR2 provide a solid base for interpreting genomic Illumina short-read sequences deriving from whole genome sequencing experiments. The allele frequencies of rDNA SNPs and InDels derived from short-read Illumina sequencing of the BACs can be related to the allele frequencies derived from short-read sequencing of the whole genome of *Arabidopsis*. This allows attributing SNPs and InDels specifically to one NOR and determine the allelic proportion for those that occur in both. Polymorphisms found on both NOR2 and NOR4, at similar frequency, would have the same allelic frequency in the BAC based Illumina sequencing (NOR2) and in the whole genome sequencing (plant). Whereas, polymorphisms more frequent on NOR2 than NOR4 have a higher allelic frequency in the BAC based Illumina sequencing than in the whole genome sequencing and *vice versa*. The largest fraction of SNPs/InDels are present within the non-coding regions of the rDNA units as previously shown ^7^ with most variations present on NOR4 (plant) (Fig 3a). One variation which is prominently enriched on NOR2 is the CAT insertion at position 4466 with ~ 9% AF in plant derived sequences and ~ 25% in the BAC derived sequences (representing only NOR2) (Fig 3b) (Table S3). Our results confirm the previous observation made by Chandrasekhara et al. 2016 ^9^. We could detect that SNPs within the 25S rDNA are mainly located on NOR4 (Figure 4d) (Table S3) in contrast to SNPs/InDels within the 18S which are located on NOR2 (Fig. 4c) (Table S3). The differential distribution of SNP/InDels between the two NORs and the higher order organization (clustering) of the rDNA units led us to design a specific fluorescence *in situ* hybridization (FISH) probe that would predominantly mark NOR2.

**Figure 4:**
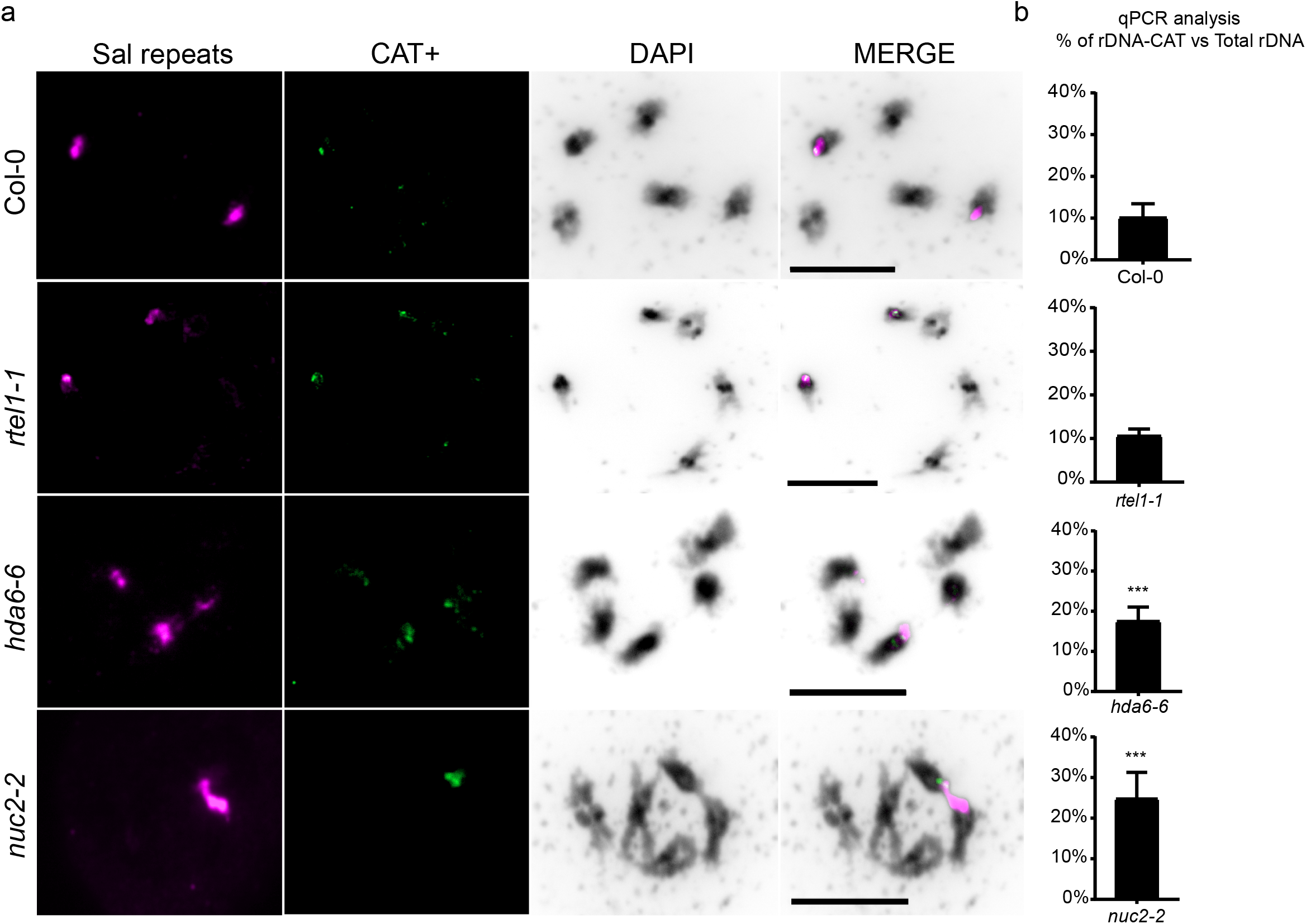
Visualizing the rDNA-CAT cluster on NOR2. **a**, Spread nuclei from pollen mother cells at diakinesis or metaphase I stages from wild type (Col-0) and mutants lines (*rtel1-1*, *hda6-6*, *nuc2-2*) hybridized for the *Sal*I repeats (magenta), the CAT insertion (CAT+, green) and stained with DAPI (black). **b**, Graphs of qPCR experiments from wild type (Col-0) and indicated mutants lines depicting the percentage of rDNA repeats bearing the CAT insertion related to the total amount of rDNA repeats. There is a statistically significant (p≤0.001) increase in rDNA-CAT copy number in *hda6-6* and *nuc2-2* mutant lines. Statistical analysis was performed using the Mann-Whitney test.

### Visualizing the CAT cluster on NOR2

In order to prove the existence of the rDNA unit sub-clusters we designed a fluorescence *in situ* hybridization (FISH) experiment that targets and marks a specific rDNA sub-cluster on condensed (meiotic) chromosomes. One of the most prominent sequence features of the rDNA units located on NOR2 is a characteristic “CAT” insertion (Fig. 3b). Our analysis suggests that rDNA units encoding this variant are clustered (Fig. 2). We anticipated that this clustering can be visualized in plant cells. We designed an LNA probe (CAT+) that specifically detects the CAT insertion located within NOR2. We performed a FISH experiment on spreads of pollen mother cells (PMCs) also employing an LNA probe directed against a universal rDNA sequence (*Sal*I probe), detecting all 45S rDNA in *Arabidopsis* ^19^ (Fig. 4a). We observed a single dominant CAT+ signal on NOR2 colocalizing with the universal *Sal*I probe in wild-type plants (ecotype Col-0), strongly supporting the results of our sequencing efforts. We also investigated mutant plants with known instability of the NORs: *rtel1-1, nuc2-2* and *hda6-6* mutants ^19–21^. In the *rtel1-1, nuc2-2* and *hda6-6* mutants we detected a different localization of the CAT+ probe. The *hda6-6* and *rtel1-1* show a redistribution of the CAT cluster to both NORs indicating a translocation event, whereas in the *nuc2-2* the CAT+ probe is much more prominent and covers a much larger area than the control.

We also developed a specific qPCR assay that allows quantification of rDNA units containing the CAT sequence and puts them in relation to the total amount of rDNA units. According to the qPCR quantification, 9% of all rDNA units contain the CAT sequence in wild-type (Col-0) plants. This is in line with the 9% allele frequency for the CAT sequence feature that we established by Illumina sequencing (Fig. 4b). We also quantified the rDNA units containing the CAT sequence in the mutant lines with NOR instability. We detect a two-fold increase of rDNA containing the CAT sequence in the *hda6-6*, *nuc2-2* mutant lines which confirms the results obtain by the *in situ* hybridization experiments (Fig. 4b). In contrast, the *rtel1-1* mutant line shows no significant difference in copy number of the CAT, although cytologically the CAT is equally distributed between the NORs. Our results confirm the presence of the rDNA-CAT sub-cluster on NOR2 and provide a tool to visualize and analyze genomic rearrangements events between the NORs.

### Tissue-specific expression and ribosomal integration of rRNA variants

The generation of the first map of NOR2 and its information regarding sequence variations within the rDNA units provide the possibility to attribute rRNAs to specific NORs and rDNA units. In this sense, we could assess whether different tissues preferentially express rDNA unit types which contribute to a heterogeneous ribosome population.

Multiple SNPs and InDels were located in the 18S and 25S rRNA genes. In order to analyze if these rRNAs variants are transcribed and integrated into ribosomes we sequenced total RNA and ribosomal RNA from different tissues: adult leaves (AL), young leaves (YL), inflorescences (INFLO) and siliques (S). To enrich for rRNAs incorporated into ribosomes we UV-crosslinked plant material and subsequently captured ribosomes with magnetic beads coupled to an LNA probe directed against the 25S rRNA. The enrichment of ribosomal rRNA was validated by RT-qPCR (tissue from adult leaves) comparing non-crosslinked and crosslinked plant material incubated with the 25S LNA probe with an unrelated control LNA probe. We find an 8-fold enrichment of ribosomal rRNA in the cross-linked sample incubated with the 25S rRNA LNA probe, compared to the control samples (Figure 5a-b).

**Figure 5:**
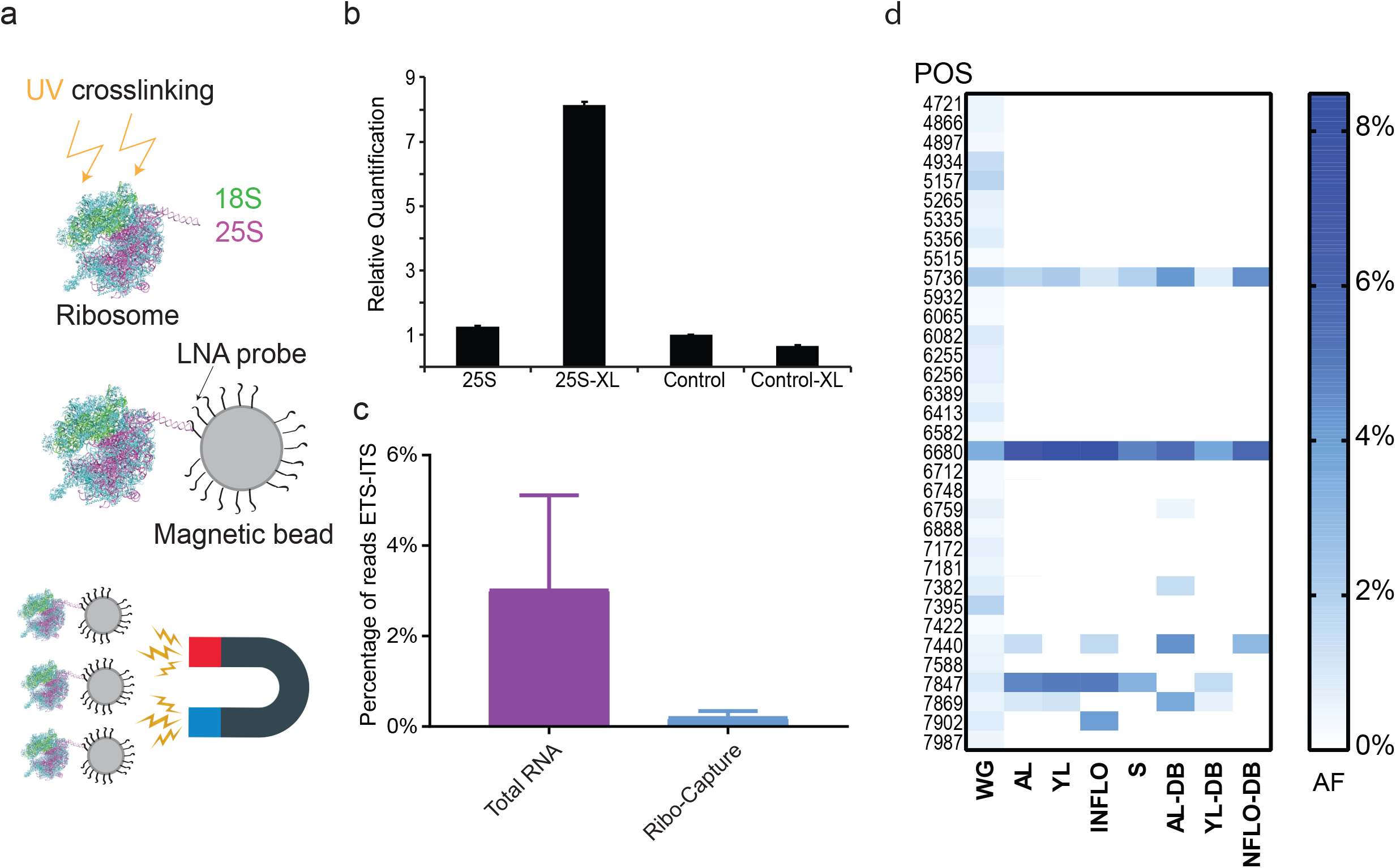
rRNA variants are differentially expressed between tissues. **a**, Ribosome capture strategy for enriching mature ribosomes. Ribosomes are UV crosslinked and captured with a magnetic bead coupled to a DNA/LNA mixamer with high affinity for the 25S rRNA. **b**, Graph depicting the relative quantification of 25S rRNA by qPCR after ribosome-capture. Non-crosslinked ribosomes captured with the 25S LNA (25S), crosslinked ribosomes captured with the 25S probe (25S-XL), non-crosslinked ribosomes captured with an LNA probe that does not bind to the ribosome (Control) and crosslinked ribosomes captured with an LNA probe that does not bind to the ribosome (Control-XL). **c**, Graph depicting the percentage of Illumina reads mapping to the ETS-ITS regions of the rDNA relative to the total amount of reads that map to the entire rDNA unit from total RNA and ribo-capture experiments. The reads from all tissues analyzed were used for this analysis. The ETS-ITS regions of the rDNA are present in the pre-rRNA but not present in mature ribosomes. **d**, Heat map representing the allele frequency of SNPs found in the 25S of mature ribosomes in different tissue types. Blue boxes represent the highest value (8%), white boxes represent the lowest value (0%). The y axis represents the positions of the SNPs with respect to the reference rDNA and are as follows: from top to bottom 4721, 4866, 4897, 4934, 5157, 5265, 5335, 5356, 5515, 5736, 5932, 6065, 6082, 6255, 6256, 6389, 6413, 6582, 6680, 6712, 6748, 6759, 6888, 7172, 7181, 7382, 7395, 7422, 7440, 7588, 7847, 7869, 7902, 7987. The x axis defines the different tissue types and sources: adult leaves (AL), young leaves (YL), inflorescences (INFLO), siliques (S), online repository derived reads (DB), DNA (whole genome sequencing data set).

The crosslinked samples, derived from 4 different tissues, enriched for ribosomal rRNAs were further processed and sequenced (Illumina short-read 150bp pair end sequencing). As a comparison, total RNA of all tissues was also sequenced in the same manner. We evaluated sequencing reads from all ribosomal-enriched and all total RNA samples that mapped to the rRNA external and internal transcribed spacer (ETS/ITS) regions in relation to the total amount of reads mapping to the reference rDNA. The ETS/ITS regions are absent in the rRNAs integrated in the mature ribosomes but present in the pre-rRNAs ^4^. We find a 10-fold lower relative enrichment of ETS/ITS reads from ribosomal-enriched material compared to total RNA, indicating that the ribosomal enrichment strategy has been successful (Figure 5c). In order to identify whether there are tissue-specific ribosomal rRNA variants, we mapped reads obtained from the ribosomal rRNA samples (cross-linked, 25S rRNA LNA probe) to the reference rDNA, filtered the reads for quality (q>30) and called SNPs and InDels with the software LoFreq. Only variations found both in the rRNA and in the rDNA that are of the same quality were taken into consideration (Table S5). Other sequence variations present in the rRNA reads which originate from secondary modifications of the rRNA were omitted from the analysis. 22% of the SNPs/InDels found in the 25S rDNA are present within the 25S rRNAs and incorporated in ribosomes (Fig. 6d, Table S5). Reads corresponding to the two SNPs at positions 5736 and 6680 within the 25S rRNA were found in all tissues but with different frequencies (Fig 5d, Table S5). It is interesting to note that, for instance, SNP 6680, which is a C to T transition, was found to have an allele frequency of 3.4% when assessing the whole genome, yet a relative incidence of up to 8.4% when analysing rRNA of mature ribosomes. Similarly, for several SNPs with low allele frequency in the whole genome (WG), corresponding reads of rRNAs were found with significantly higher frequency in mature ribosomes: SNP at position 7440 (WG=0.5% AF) is expressed in adult leaves (1.4% AF) and inflorescences (1.6% AF); SNP at position 7869 (WG= 0.57% AF) is expressed in adult (1% AF) and young leaves (1.2 % AF) (Fig. 5d) (Table S5). We could not detect reads corresponding to SNPs or InDels in the 18S rDNA in samples enriched for rRNA integrated into ribosomes.

To corroborate our findings, we analyzed previously published data sets of polysome profiling experiments performed in *A. thaliana* Col-0 plants using the same tissues indicated above ^22–25^. Polysomes are the cellular fraction of ribosomes which are actively translating single mRNAs. While the focus of these previous studies was mRNA, the data sets contained a large number of rRNA-related reads that we mapped to the reference rDNA ^22–25^. More than 60% of the SNPs identified in our ribosome capture assays where also found in the polysome datasets (Table S5) (Fig. 5d).

This set of experiments confirms that the rRNA gene variants we detect in the rDNA are expressed in a tissue-specific manner and present in translating ribosomes. Furthermore, the presence of only 25S rRNA variants suggests that NOR4 is mainly contributing to the pool of ribosomes even within tissues that have both NOR2 and NOR4 transcriptionally active (Young Leaves and Inflorescences)^6,19^.

## Discussion

We have generated a comprehensive assembly of the *Arabidopsis thaliana* nucleolus organizer region of chromosome 2 by using a BAC based approach ^12^. Embedded in our study we provide a sequencing and bioinformatic pipeline with *ad hoc* generated programs to aid the assembly of large and complex repetitive regions and address their higher order organization.

The assembly and analysis of the NOR2 contigs allowed us to draw conclusions on the overall higher-order organization of the rDNA units and their heterogeneity. Our data show that the rDNA units are heterogeneous with many large and small variations present in the internal and external transcribed spacer regions and, to some extent, in the core rRNA subunits. While some of the sequence variants have already been identified before ^7,9^ our data puts all of them in context of large sequence contigs. Our draft assembly is sufficiently complete to demonstrate that there are specific rDNA clusters that share similar sequence features which could have originated from former rDNA homogenization events ^3^. These events have led to the formation of large regions that contain rDNA units of the same type (100 kb, 10 rDNA units). Furthermore, our data allows attribution of processed rRNAs to their origin from chromosome 2 or 4. Previously, only rDNA and unprocessed rRNA could be distinguished and attributed according to few polymorphism and their 3’ ETS variant ^6,8,26^.

Based on a comparative analysis of short-read sequencing experiments performed on BACs (exclusively from NOR2) and genomic DNA we were in the position to determine which SNPs/InDels are enriched on NOR2 and which on NOR4. Most of the unique sequence variants appear to be present on NOR4 or shared between NOR2 and NOR4 exception made for the CAT sequence insertion which is present only on NOR2.

The presence of rDNA clusters is supported by the analysis of spread meiotic chromosomes. Utilizing an LNA probe specifically directed against the CAT insertion rDNA unit type we detected signals only on NOR2. In addition, the probe allowed us to visualize rDNA expansion and re-arrangement events in the *hda6-6*, *nuc2-2* and *rtel1-1* mutant lines which are known to have unstable rDNA copy numbers ^19,20^. The expansions were further corroborated by qPCR experiments. We herewith generated the information and the tools to analyze (meiotic) recombination within and between the repetitive and highly similar rDNA domains by fluorescence *in situ* hybridization.

Interestingly, only up to 20% of the SNPs contained within the 25S rDNA were detected in mature ribosomes. The two SNPs at positions 5736 and 6680 which are expressed in all tissues analyzed, are located within the ribosomal extension segment (ES) 12 and the ES27 of the large ribosomal subunit which are known to be highly variable regions ^27^. In contrast, the SNP at position 7440 which is expressed only in adult leaves and inflorescences is located on the loop between H74 and H88 which is located in the ribosome peptidyl transferase site^28^. This SNP is a G to T transition and could have a functional impact on the ribosome by changing the interaction of the tRNA with the ribosome.

Our data implies that only a minor subset of the rRNA genes is transcribed at a given time in a well-controlled manner. Furthermore, none of the larger deletions or insertions that were detected in rDNA units (Suppl. Fig. 3b-c) are detected in the total RNA datasets or in the mature ribosomes. This could indicate that these large deletions perturb the maturation of the rRNA into fully folded ribosomes. It is also interesting to note that we could never detect rRNAs bearing the CAT insertion in any of our and previously published ^22–25^ total RNA sequencing datasets. This indicates that even though both NORs are transcriptionally active in young seedlings ^29^ and in inflorescences ^19^, not all repeats of NOR2 are transcribed. This is in contrast to what was previously hypothesized since NOR2 is not transcriptionally silent in adult somatic tissues. These results reveal a complex, multi-level regulation of rRNA gene transcription and ribosomal rRNA integration. The locus-specific epigenetic silencing of NOR2, as a whole, was previously shown only on the base of the expression of the 3’ETS ^9^. In contrast, our datasets, with the precise positioning of each rDNA unit to a defined cluster, suggest that rDNA silencing might be confined to the clusters themselves. We conclude that clusters of rRNA unit types are co-regulated in a tissue-specific manner supporting the concept of tissue-specific ribosome subpopulations differing in their rRNA composition.

## MATERIALS AND METHODS

### BAC library

The Bacterial Artificial Chromosome library used in this study was established by Mozo et al. in 1998 ^14^ at the “Institut für Genbiologische Forschung” (IGF) in Berlin, Germany. The library is available at the Arabidopsis Biological Resource Center (ABRC) (https://abrc.osu.edu/).

### BAC extraction

BAC DNA extraction has been carried out according to the “Qiagen Large Construct extraction protocol” (April, 2012) which has been specifically designed to retrieve large amounts of long, circular and intact DNA, such as BACs and Cosmids.

A bacterial culture or a single colony was inoculated in 500 ml of LB liquid media containing the appropriate selective antibiotic (kanamycin, final concentration 100 μg/ml) and grown overnight at 37 °C with vigorous shaking (300 rpm). Cells were harvested through centrifugation (6000xg) for 15 minutes at 4 °C. 6000g corresponds to 6000 rpm in Sorvall GSA or GS3 of Beckman JA-10 rotors. The pellet was resuspended in 20 ml of Buffer P1 (Resuspension Buffer) containing RNase A (with final concentration 100 μg/ml). The cells were lysed by adding 20 ml of Buffer P2 (Lysis Buffer). pH equilibration was achieved by adding 20 ml of Buffer P3 (Neutralization Buffer). After centrifugation at >20000xg for 30 minutes at 4°C (a centrifugal force of 20000xg corresponds to 12000 rpm in a Becman JA-17 rotor or 13000 rpm in a Sorvall SS-34 rotor), the supernatant containing the BAC DNA was transferred to a new centrifugation tube. A cell strainer (70 μm) was used in order to separate effectively cell debris from the supernatant. The DNA was precipitated by adding 35 ml of isopropanol at room temperature and collected by centrifugation (>15000xg for 30 minutes at 4°C) and washed with 5 ml of ethanol (70%). After removal of alcohol the pellet was air-dried for 10 minutes.

To select for circular and unfragmented DNA, an exonuclease digestion step was performed (NEB T5 exonuclease, 10000 unit/ml). 9.5 ml of Reaction buffer were added to the dried pellet for resuspension, followed by addition of 500 μl of EX buffer and 1 μl of exonuclease and incubation at 37°C for 1 hour. In the meantime, the filtration columns were equilibrated (QIAGEN-tip 100) with 15 ml of buffer QBT (Equilibration Buffer). Then 10 ml of Buffer QS (Adjustment Buffer) were added to the sample and the sample was thereafter loaded onto the column. After the DNA has entered the resin, 60 ml of buffer QC (Wash buffer) were added. The BAC DNA was eluted by adding 15 ml of Buffer QF (Elution Buffer).

The DNA was precipitated by the adding isopropanol (as above) and then washed once with 70% ethanol. After having removed the ethanol, the pellet was air-dried for 20 minutes and then resuspend in 100 μl of TE buffer. To enhance DNA resuspension, TE buffer was pre-warmed to 65 °C.

### BAC linearization

BACs extracted from *E. coli* were linearized with the restriction enzyme *Apa*I. This enzyme was selected since it is a single cutter in the vector backbone (pBeloBAC-Kan) and according to available sequence information leaves the rDNA repeats intact. The reaction was incubated overnight at 37 °C. Enzyme inactivation was carried out by incubating the sample at 65 °C for 20 minutes.

### Pulse Field Gel Electrophoresis

The successful linearization of the BACs has been assessed through Pulse Field Gel Electrophoresis (PFGE) using the Clamped Homogeneous Electric Fields (CHEF) technology. 2% low melting agarose solution (Certified™ Low Melt Agarose, BioRad) was prepared using 1X TBE buffer (Tris base 1M, Boric Acid 1M, EDTA 0,02M) and melted using a microwave. The solution was equilibrated at 50°C in a water bath. 1 μg of linearized BAC DNA was combined with equal volume of 2% Certified™ Low Melt Agarose and mixed by pipetting in order to achieve a final concentration of 1% agarose. The mixture was quickly transferred to plug molds and left to solidify for 10 minutes. While waiting for the solidification of the plugs, 1% agarose gel (Pulse Field Certified Agarose, BioRad) was prepared using 1X TBE buffer and casted in a PFGE casting stand. Finally, the plugs were inserted inside the wells. PFGE was run according to the following program for 20 hours: Switch time 200 seconds, Reorientation Angle 120 °C, Voltage gradient 5V/cm. The final result was visualized through a transilluminator by staining the gel with EtBr for 1 hour in 1X TBE.

### rDNA-CAT copy number quantification

DNA was extracted by crushing leaves in UREA buffer (0.3 M NaCl, 30 mM TRIS-Cl, pH 8, 20 mM EDTA, pH 8, 1% [w/v] N-lauroylsarcosine, 7 M urea) and subsequently purified with phenol:chloroform:isoamylalcohol (25:24:1). The qPCR reaction was performed using the KAPA SYBR FAST kit following the product specifications. To quantify rDNA-CAT copy numbers, 20–30 ng of genomic DNA were used together with two primer pair sets: 18SRealdn and 18SRealup as previously described for *Arabidopsis* (Muchová et al., 2015) to amplify all rDNA copies, CAT+ fw (CGC ATC AGC AAA GGA TGA TGG) – CAT+ rv (AGT CTA AAA CGA CTC TCG GCA) to amplify only the rDNA-CAT. As a Ct calibrator for the experiment BAC F2J3 was used since it has 9 rDNA copies of which one with the CAT insertion. Additionally, for each genomic sample, the reference gene ACTIN-7 was used to calculate the rDNA copy number. The amplification of the rDNA, rDNA-CAT and of the reference gene ACTIN-7 was performed in separate wells and in three technical replicates. The analysis was conducted with two separate biological replicates. The conditions used for the qPCR were 95°C 1 min initial denaturation, 95°C 30 s, 65°C 30 s, 72°C 30 s for 40 cycles with fluorescence detection after every elongation step. The PCR products were not longer than 250 bp and contained a GC content of ∼50%. The experiment was performed on an Eppendorf Realplex 2 Mastercycler.

### 3’ETS variant PCR

Amplification of the 3’ETS variants was performed as in Pontvianne et al. ^19,30^

### Westburg Preparation for Illumina sequencing

BACs were prepared for Illumina sequencing according to the “Westburg NGS DNA library prep kit” (protocol 3.1 10/2018) with minor optimizations. Firstly, the DNA was fragmented through an enzyme mix provided in the kit. 5 μl of fragmentation mix (10X) are added to 100 ng of DNA (final volume of the reaction: 40 μl) and left for 30 minutes at 65 °C, after an equilibration period at 32 °C for 20 minutes.

Secondly, the DNA fragments are linked to the sequencing adaptors and barcoded through PCR. Illumina adaptor are mixed together equimolar amounts of forward and reverse oligos to obtain a final concentration of 1.5 μM. The mix was incubated at 95°C for 5 minutes on a thermocycler, and left to cool down for 30 minutes. 2.5 μl of the annealed Illumina adapter is mixed with 40 μl of the fragmented DNA together with 17.5 μl of H_2_O, 20 μl 5X ligation buffer and 10 μl of DNA ligase. Samples were incubated at room temperature for 15 minutes. 80 μl of AmpureXP beads were added and mixed by pipetting or vortexing. Captured magnetic beads are washed once with 85 % EtOH and resuspended in 50 μl of water. 20 μl of Binding solution are added (20 % PEG8000, 2.5M NaCl, 0.05 % Tween20), mixed thoroughly and incubated 5 minutes at room temperature. After capturing the beads the supernatant is transferred to a fresh well and 20 μl of new beads added. The samples are incubated for 5 minutes at room temperature. The beads were washed twice with 85 % ethanol and left to dry for 1 minute. The samples were suspended in 11 μl of water for 2 minutes at room temperature. After capturing the beads 10 μl of the supernatant were transferred to a fresh well. 15 μl of 2x KAPA HF Ready Mix were added with 2 μl of the appropriate barcode, 2μl of the TrueSeq Universal Adapter and 1 μl of H_2_O. The sampels were ran with the following PCR program:

| Step | Temperature | Time |
| --- | --- | --- |
| DNA polymerase activation | 98 °C | 1 minute |
| 15 X | 98 °C | 5 seconds |
|  | 65 °C | 30 seconds |
|  | 72 °C | 1 minute |
| Final elongation | 72 °C | 1 minute |
| Hold | 4 °C |  |

Finally, the samples were purified and cleaned by Ampure XP beads. 27 μl of Ampure XP beads were mixed by pipetting or vortexing with the sample. The beads were captured and the supernatant was discarded; the pellet was washed twice with 85% ethanol. The beads were resuspended in 25 μl of water, the beads were captured and the supernatant transferred to a fresh well. The samples were now submitted for sequencing.

### Nanopore library preparation with native barcoding

| Protocol Step | Reagent | Kit name or code |
| --- | --- | --- |
| DNA repair and end-prep | NEBNext FFPE DNA Repair Buffer | M6630 |
|  | NEBNext FFPE DNA Repair Mix |  |
|  | Ultra II End-prep reaction buffer | NEB E7645 |
|  | Ultra II End-prep enzyme mix |  |
| Barcode Ligation | Native barcodes (NBD01 -NBD12) | Native Barcoding Expansion 1-12 (EXP-NBD104) |
|  | NEB Blunt/TA Ligase Master Mix | M0367 |
| Adapter ligation and clean-up | Adapter Mix II (AMII) | Native Barcoding Expansion 1-12 (EXP-NBD104) |
|  | NEBNext Quick Ligation Reaction Buffer (5X) | NEB B6058 |
|  | Quick T4 DNA Ligase 2M U/ml | NEB M0202 |
|  | Long Fragment Buffer (LFB) | Ligation Sequencing Kit (SQK-LSK109) |
|  | Elution Buffer (EB) | Ligation Sequencing Kit (SQK-LSK109) |
| Priming and loading the SpotON flow cell | Sequencing Buffer | Ligation Sequencing Kit (SQK-LSK109) |
|  | Flush Tether (FLT) | Ligation Sequencing Kit (SQK-LSK109) |
|  | Flush Buffer (FB) | Ligation Sequencing Kit (SQK-LSK109) |
|  | Loading Beads (LB) | Ligation Sequencing Kit (SQK-LSK109) |

Linearized BACs have been sequenced through Oxford Nanopore Technology. Twelve linearized BACs were multiplexed, exception made for BACs F2J17 and F1E12 which were processed individually (Table S1), and sequenced with a Nanopore MinION using an R9.4 flowcell (FLO-MIN109, ONT). For each BAC only ultra-long reads (longer than 50 kb) were used for the assembly, exception made for BAC F1L21 (Suppl. Fig 1c.) (Table S1). Each BAC was assembled by using the software CANU ^31^ and polished three times using the software Nanopolish ^32^.

Multiplexing preparation followed the manufacture’s manual “1D Native barcoding genomic DNA -SQK-LSK109” (version: NBE_9065_v109_revH_23May2018) with some optimizations.

The protocol is divided into 4 main steps:

- DNA repair and end-prep
- Barcode ligation
- Adapter ligation and clean-up
- Priming and loading the SpotON flow cell

#### DNA repair and end-prep

In the first step, DNA repair was achieved through the utilization of a cocktail of enzymes (NEBNext FFPE DNA repair Mix) designed specifically to repair breaks in the DNA. This cocktail is able to repair nicks, gaps (up to 10 nucleotides), blocked 3’ ends, oxidized bases and deamination of cytosine to uracil. Afterwards, the DNA ends are repaired through the Ultra II End-prep enzyme mix, which enhances the attachment of the DNA barcodes. It is important to note that BACs are extremely sensible to physical sharing. For this reason, throughout this protocol, we mixed the samples always by manual flicking to avoid to pipette and vortex. 1.2 – 1.5 μg of DNA were transferred into a 1.5 ml DNA LoBind tube and the volume adjusted to 48 μl with Nuclease-free water and mixed thoroughly by inversion or flicking. We added to the DNA following exactly this order directly in the LoBind tube, 3.5 μl NEBNext FFPE DNA Repair Buffer, 2 μl NEBNext FFPE DNA Repair Mix, 3.5 μl Ultra II End-prep reaction buffer and 3 μl Ultra II End-prep enzyme mix. The samples were incubated at 20 °C for 5 minutes and 65 °C for 5 minutes. During the incubation we prepared 500 μl of fresh 80 % ethanol in Nuclease-free water. We added 60 μl of resuspended AMPure XP beads to the end-prep reaction and mixed by flicking the tube. The samples were incubated on a Hula mixer (rotator mix) for 15 minutes at room temperature, then spun down briefly and placed on a magnetic rack. We kept the LoBind tube on the rack and pipetted off the supernatant, then the beads were washed twice with 200 μl of 80 % ethanol. The samples were spanned down and placed back on the magnet. The residual ethanol was removed and the samples left to air dry for 60 seconds. The samples were resuspend by flicking in 25 μl Nuclease-free water. To enhance elution, we incubated the samples for 20 minutes at 37 °C by using a thermocycler.

The pellet was spun down and the sample placed on a magnetic rack until the eluate is clear and colorless, we removed and retained 25 μl of eluate into a clean 1.5 Eppendor DNA LoBind tube.

1 μl of end-prepped DNA was measured using a Qubit fluorometer (recovery aim > 200 ng). We then proceeded with the naïve barcoding step.

#### Barcode ligation

The second step consists on barcoding the samples in order to sequence them together. We added to the 24 μl of repaired and end-prepped DNA, 2.5 μl Native Barcode (Native Barcode 1-12, EXP-NBD104) and 25 μl Blunt/TA Ligase Master Mix, maintaining this order. We mixed by flicking and briefly spanned down. The samples were incubated at room temperature for 10 minutes. We added 50 μl of resuspended AMPure XP beads to the reaction and mixed by flicking, then incubated the sample on a rotator mix for 15 minutes at room temperature. The sample was spanned down and the sample placed on a magnetic rack; the supernatant was removed and the beads washed twice with 80 % ethanol. Residual ethanol was removed and the sample left to air dry for 60 seconds (do not dry the pellet to the point of cracking). The tube was removed from the magnetic rack and resuspended by flicking in 26 μl Nuclease-free water. To enhance elution, the sample was incubated for 20 minutes at 37 °C. The sample was palced on a magnetic rack until the eluate was clear and colorless. We then removed and retained 26 μl of eluate containing the DNA library into a clean 1.5 Eppendor DNA LoBind tube.

The barcoded samples were pooled together into a 1.5 ml Eppendorf DNA LoBind tube, ensuring that sufficient sample is combined to produce a pooled sample of > 400 ng total. It is not fundamental to pool equimolar amounts of the samples, as long as the concentrations are not too different (differences in the final quantity of barcoded samples between 50 and 80 ng are acceptable). If the volume of the pooled samples exceeded 65 μl, it was necessary to perform an additional concentration step.

Pooled barcoded samples were quantified using a Qubit fluorometer (recovery aim >500 ng).

#### Adapter ligation and clean-up

The final step consists on the ligation of the adapters needed for sequencing to the DNA fragments. This is achieved by adding the proprietary Nanopore adaptor (Adapter Mix II) with the barcoded DNA and the T4 DNA ligase. Adapter ligation was performed by mixing by flicking the tube between each sequential addition of: 65 μl pooled barcoded sample, 5 μl Adapter Mix (AMII), 20 μl NEBNext Quick Ligation Reaction Buffer (5X) and 10 μl Quick T4 DNA Ligase.

The samples were mixed by flicking incubated for 10 minutes at room temperature. 50 μl of resuspended AMPure XP beads were added to the reaction and mixed by flicking, then incubated on a rotator mixer for 15 minutes at room temperature. The samples were placed on a magnetic rack and the beads allowed to pellet. The supernatant was removed and the beads washed twice by adding 250 μl Long Fragment Buffer (LFB) and left to dry for 30 seconds. The sample was resuspended in 15 μl Elution Buffer (EB) and incubated for 20 minutes at 37°C. The samples was placed on a magnetic rack until the eluate is clear and colorless. The supernatant was removed and retained into a clean 1.5 ml Eppendorf DNA LoBind tube. The library was now ready to be sequenced.

#### Priming and loading the SpotON flowcell

In this step the flow cell is put under pressure and prepared for sequencing by the addition of the appropriate buffers. Pressurization enhances the correct flowing of the buffers in the device. The sample were mixed with the Loading Beads that take the DNA molecules close to the pores.

The flowcell was put under pressure by opening only the priming port, a P1000 pipette was set to 200 μl, inserted the tip into the priming port and turned the wheel until the dial shows 200-230 μl, or until it was possible to see a small volume of buffer entering the pipette tip. The flow cell priming mix was prepared by adding 30 μl of thawed and mixed Flush Tether (FLT) directly to the tube of thawed and mixed Flush Buffer (FB) and mixed by pipetting up and down. 800 μl of the priming mix were loaded into the flow cell via the priming port, avoiding air bubbles.

The final library was prepared for sequencing by mixing together 37.5 μl Sequencing Buffer (SQB) with 25.5 μl Loading Beads (LB) (taking care to mix them immediately before use since they precipitate quickly) and 14 μl of the DNA library. The prepared library was mixed by gently pipetting up and down just prior to loading. We completed the flow cell priming by gently lifting the SpotON sample port cover to make the SpotON sample port accessible, then by loading 200 μl of the priming mix into the flow cell via the priming port, avoiding the introduction of air bubbles. The prepared DNA library was mixed gently and loaded to the flow cell via the SpotON sample port in a dropwise fashion. The sequencing runs were carried out until all pores were completely exhausted and generated on average 7.12 Gb of raw sequence information (Tables S1, S2).

### *Arabidopsis thaliana* growth conditions

*Arabidopsis thaliana* Col-0 ecotype seeds were stratified in water in the dark at 4°C for two days before sowing on soil/perlite 3:1 mixture (ED 63, Premium Perlite). Pots were covered with a transparent lid until cotyledons were fully developed and first primary leaves visible. Plants were grown under long day conditions in controlled environment rooms (16 hours of light, 8 hours of darkness, 60-80% humidity, 21°C, 15,550 lux, T5 Tube illumination).

### DAPI spreads

Inflorescences were harvested into fresh fixative (3:1 96% [v/v] ethanol [Merck] and glacial acetic acid) and kept overnight (O/N) for fixation. Once the fixative decolorized the inflorescences, they were placed in fresh fixative (can be stored for over a month at −20°C), and subsequently, one inflorescence was transferred to a watch glass. The yellow buds were removed to collect only white and transparent buds. The white buds were separated from the inflorescence and grouped according to size. This step is necessary to obtain preparations with separated meiotic stages.

Afterward, the buds were washed three times with citrate buffer (0.455 mL of 0.1 M citric acid, 0.555 mL of 0.1 M trisodium citrate in 10 mL of distilled water) and submerged in an enzyme mix (0.3% w/v cellulase, 0.3% w/v pectolyase in citrate buffer). Each bud has to be submerged for the digestion to work efficiently. The buds were incubated for 90 min in a moisture chamber at 37°C. Digestion was inhibited by adding cold citrate buffer. At this point, the buds were transferred (maximum three to four buds of the same size) to a glass slide. Excess liquid was removed and 15 μl of 60% acetic acid added. The buds were suspended using a metal rod and an additional 10 μl of 60% acetic acid was added to the suspension. The droplet area was labeled using a diamond needle and fixed with fixative. Slides were dried for at least 2 h. To stage the meiocytes, 15 μl of 2 μg/ml 4’,6 diamidino-2-phenylindol (DAPI) diluted in Vectashield (Vector Laboratories) was added to the slide and sealed with a glass cover slip. Images were taken on a Zeiss Axioplan microscope (Carl Zeiss) equipped with a mono cool-view charge-coupled device camera ^33^.

### Fluorescence *in situ* hybridization

The slides selected for fluorescence *in situ* hybridization (FISH) were washed with 2x SSC for 10 minutes and incubated for 2 minutes at 37 °C in pre-heated 0.01M HCl with 250 μl of 10 mg/ml of Pepsin and 5 μl of 100 mg/ml RNAseA. The slides were then washed in 2x SCC for 10 min at room temperature. Fifteen microliters of 4% paraformaldehyde were added onto the slides, covered with a strip of autoclave bag, and placed for 10 min in the dark at room temperature. The slides were then washed with deionized water for 1 min and dehydrated by passing through an alcohol series of 70, 90, and 100%, for 2 min each. Slides were left to air dry for 30 min.

Meanwhile, the probe mix was prepared by diluting 1 μL of probe (2–3 μg of DNA) in a total of 20 μL of hybridization mix (10% dextran sulfate Molecular Weight [MW] 50,000, 50% formamide in 2× Saline Sodium-Citrate [SSC]).

In case the rDNA Locked Nucleic Acids (LNA) probe was applied, only 50 pmol (final concentration) was used per slide. The probe mix was denatured at 95°C for 10 min and then placed on ice for 5 min. Afterwards, the probe mix was added to the slide, covered with a glass cover slip, sealed, and placed on a hot plate for 4 min in the dark at 75°C. Finally, the slides were placed in a humidity chamber overnight at 37°C. After hybridization, the cover slips were carefully removed and the slides were treated with 50% formamide in 2× SCC for 5 min in the dark at 42°C. The slides were then washed twice with 2× SCC for 5 min in the dark at room temperature.Slides were visualized on an inverted epi-fluorescent microscope equipped with an CCD camera. LNA probes used in this study are: ATG+ (CCC TCA CCA TCA TCC TT) labelled with TYE563 (red) and *Sal*I ^19^.

### Cleaved Amplified Polymorphic (CAPs) Marker confirmation

1 μg of DNA from BACs F2D9, F2G3, F2G18, F2E13, F2I6 were tested for the presence or absence of the restriction site (*Ava*I) at position 4133 of the reference rDNA. The region harboring the restriction site was amplified by PCR (primer sequences: AVAI Fw CGCATCAGCAAAGGATGATGG, AVAI Rv ACCTTGGGATGGGTCGG). The final amplicons were digested with the enzyme *Ava*I. 17 μl of the PCR reaction were mixed with 2 μl of CutSmart Buffer and 1 μl of *Ava*I (NEB) for a final volume of 20 μl. The reaction was incubated at 37 °C overnight. The samples were loaded on an agarose gel (1 % w/v + EtBr) and visualized on a Biorad transilluminator.

### Ribo-Capture Protocol

Ribosome captures were performed as in Rogell et al. ^34^ with some modifications. Magnetic beads (Dynabeads M-270 Carboxylic Acid) were activated and coated with *ad hoc* designed LNA probes (Qiagen), harboring an −NH_2_ group, following the manufacturer’s instructions. The beads were stored for a maximum of 3 months. Ribosome capture was performed as explained in Rogell et al. In brief, plants were UV (260 nm) crosslinked in a Stratalinker at 0,15 J/cm^3^ and placed at 10 cm from the light source. After irradiation the plant material was immediately frozen in liquid nitrogen and stored at −80 °C. Plant material was ground in 1 ml Lysis Buffer (20 mM Tris-HCl [pH 7.5], 500 mM LiCl, 0.5% LiDS [w/vol, stock 10%], 1 mM EDTA and 5 mM DTT) with the addition of 1μl of Protease Inhibitor (Thermo Fisher) and 1 μl of RNAse inhibitor (RiboLock). 15 % v/v of formamide or ethylene carbonate were added to the sample as a hybridization enhancer. The samples were placed at 60 °C for 5 min. Beads coated with the LNA probe were denaturated at 95 °C for 3 min and added to the sample. The mix was placed at 40 °C for 2 h shaking. Afterwards, the samples were placed in a magnetic rack and washed 3 time with Lysis Buffer, twice with Buffer 1 (20 mM Tris-HCl [pH 7.5], 500 mM LiCl, 0.1% LiDS [wt/vol], 1 mM EDTA, and 5 mM DTT), Buffer 2 (20 mM Tris-HCl [pH 7.5], 500 mM LiCl, 1 mM EDTA, and 5 mM DTT) and Buffer 3 (20 mM Tris-HCl [pH 7.5], 200 mM LiCl, 1 mM EDTA, and 5 mM DTT). The captured RNA was eluted in 100 μl of RNAse free water for 3 min at 90 °C. Ribosomal RNA was purified by a DNAseI (Thermo Fisher) clean up, Proteinase K clean up and phenol:chloroform:isoamylalcohol (25:24:1) pH 5 extraction. At this step, the samples are ready for library preparation and RT-PCR. The Lexogen CORALL RNA-seq library kit was used for preparing the samples for Illumina sequencing. cDNA for qPCR analysis was obtained by using the iSCRIPT kit following the manufacturer’s instructions. The qPCR reactions were performed using the KAPA SYBR FAST kit following the product specifications.

25S directed LNA probe sequence: AACGCCGAAGACGTCCGAT

Mock LNA probe sequence: AAGACGTCGAAGGTTACCT

### Total RNA sequencing

Plant material from different *Arabidopsis* tissues was collected and frozen immediately in liquid nitrogen. To extract RNA, the SV Total RNA Extraction Kit (Promega) was used following the product specifications. Total RNA was further purified by a DNAseI (thermofisher) clean up and phenol:chloroform:isoamylalcohol (25:24:1) pH 5 extraction. At this step, the samples are ready for library preparation and RT-PCR. Lexogen CORALL RNA-seq library kit was used for preparing the samples for Illumina sequencing.

## BIOINFORMATIC METHODS

### MinION Sequencing

MinION sequencing was performed according to manufacturer’s guidelines using R9.4 flowcells (FLO-MIN109, ONT). The run was monitored using Oxford Nanopore Technologies MinKNOW software. The specific version of the software has changed from run to run and is included in the fast5 files generated. The base-calling has been conducted with Albacore (version 2.0) for BACs F23H14, F2J17, F1E12 and Guppy (version 2.2.3) for all the other BACs. A list of all Nanopore sequencing runs is described in Suppl. Table S8.

### Assembly

Assembly has been conducted with Canu (v1.7.1 and v1.8) starting from fastq files with reads longer than 50 kb. It is important to note that assemblies carried out by using reads with a Phred quality score greater than 9 required less reads than those with a low-quality score. The command used is the following for all the BACs:

canu overlapper=mhap utgReAlign=true -d /path/to/assembly_directory -p assembly_prefix --t 40 genomeSize=0.1m -nanopore-raw ont_reads.fastq

The MHAP algorithm was chosen because of its ability to handle repetitive elements. utgReAlign= true allows to compute a sequence alignment for each overlap produced by mhap. The final assemblies (draft_assembly.fasta) are outputted in the fasta format.

### Assembly polishing using Nanopolish

Assemblies generated by Canu have been refined using the nanopolish consensus-calling algorithm (v0.10.2 and v.0.11). Each assembly has been polished using all the reads contained in the original sequencing fastq file.

minimap2 (version 2.14) was used to map the reads to the draft assembly. The command and the options used were the following:

minimap2 -ax map-ont -t 8 draft_assembly.fasta ont_reads.fastq| samtools sort -o mapped_ont_reads.bam -T mapped_reads.tmp

nanopolish was used to produce an improved consensus sequence for the draft BAC assembly produced by Canu. The command and the options used were the following:

nanopolish variants –-consensus -o polished_assembly.vcf -r ont_reads.fastq -b mapped_ont_reads.bam -g draft_assembly.fasta

nanopolish outputs a Variant Calling Format (VCF) file (polished_assembly.vcf). To generate the polished genome in fasta format it is necessary to run the following command:

nanopolish vcf2fasta -–skip-checks -g draft_assembly.fasta polished_assembly.vcf > final_assembly.fasta

### SNPs and InDels calling

SNPs and InDels present on each BAC have been identified through Illumina sequencing. Illumina reads were mapped to the reference ribosomal DNA repeat and SNPs/InDels calling was performed using LoFreq ^16^. Reads were mapped using bowtie2 ^15^ and sorted with samtools ^35^.

The quality score for SNPs/InDels was calculated with LoFreq:

lofreq indelqual –-dindel mapped&filetered_illumina_reads.sorted.bam -f reference_repeat.fasta > final_illumina_reads.bam

SNPs and InDels were called with the following command:

lofreq call -–call-InDels final_illumina_reads.bam -f reference_repeat.fasta > called_variations.vcf

The same procedure was used for calling the SNPs/InDels in the whole genome samples (Kurzbauer et al. 2020 submitted) and the ribo-capture samples.

### rDNA reference repeat

The rDNA reference repeat (Suppl. File 2) was generated starting from one rDNA gene of BAC F2J17 (variant 1). The repeat was polished by Pilon (5 rounds) using whole genome *A. thaliana* Illumina reads (Kurzbauer et al. 2020 submitted, PRJNA555773). The *Sal*I boxes were removed from the reference.

### OverBACer (Overlap BAC finder)

To identify the overlaps between BAC assemblies, a python script was devised named OverBACer. OverBACer exploits the different lengths of SalI boxes, 3’ETS variants and other large InDels occurring within the rDNA repeats, to find the overlapping genes between two or more BACs.

OverBACer takes as input a file containing the assemblies to be analyzed (in fasta format), a search file consisting of the probes to be identified in the assemblies (in fasta format), and the maximum/minimum number of repeats needed to call an overlap.

The algorithm is divided into three steps: reference point identification, distance calculation between such points and overlap.

In the first step, OverBACer identifies the position in the assemblies of the given probes present in the search file. It accomplishes this task by performing a “word search” within the assembly. Every time a probe finds a match on the sequence, its absolute position on the assembly is annotated; these positions are called “reference points”.

In the distance calculation step, OverBACer computes the distances between neighboring reference points. This system allows to “barcode” the sequence, creating a fingerprint of each assembly. Since each rDNA repeat has a slightly different *Sal*I box length, the distances between two reference points will vary among repeats, allowing to distinguish different rDNA genes. Moreover, the combination of the distances of each repeat within an assembly assigns a unique tag to each BAC.

In the overlap step, the barcodes of each assembly are compared with all the others, thus finding the overlaps. This task is performed by checking the correct order of the reference points on the sequences and the distances associated to the reference points. In particular, a tolerance chosen by the user is applied when comparing the distances, thus taking into account the presence on the sequence of small variations such as SNPs and small InDels that might prevent the identification of correct overlaps.

Usage example:

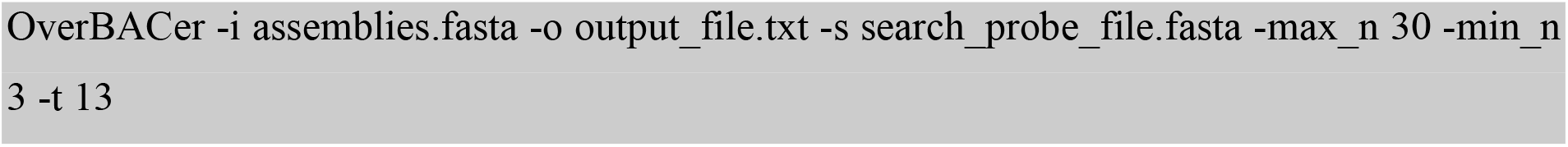

Where -i specifies the input file containing the assemblies to be overlapped, -o the output file, -s the search file consisting on the probes to be used to identify the reference points, -max_n and -min_n the maximum/minimum number of reference points to call an overlap and -t the tolerance applied when the distances are compared (in bp). The program was calibrated to find a minimum of 2 overlapping repeats with an error range of +/− 35 bp.

### Neighbour finder

To study the association of *Sal*I boxes and small SNP/InDels of rDNA repeats at 10, 20 and 30 kb, a python script (Neighbour finder) was designed.

Neighbour finder takes as input a text file consisting of the barcoded BACs classified through the alphanumerical system As explained in the results section. It analyzes the co-occurrence *Sal*I boxes (-a 0 option) and of small SNP/InDels (-a 1 option). For all pairwise combinations of barcodes, we count how often a pair of barcodes occurs at 10 kb, 20 kb and 30 kb distances. Fig 3d, e, f show the results. As an example, the association of *Sal*I boxes is described below. In both cases, it is possible to randomize the data by using the -r 1 option.

Neighbour finder extracts from the barcodes the portion of the alphanumeric code that describes the *Sal*I box length, which is represented by the letters between round brackets. Then, the code of two neighbouring repeats (at 10, 20 or 30 kb of distance) is fused to give a combination of *Sal*I boxes, which is then searched within the pool of all the combined codes generated for each BAC. Neighbour finder counts the occurrence of each combined code within the pool and returns this number associated with the corresponding code. The process is repeated three times in order to evaluate the association of *Sal*I boxes at 10, 20 and 30 kb.

Usage example:

neighbor_finder -i input_file -o output_file -t temporary file -a 0 -r 0

Where -i specifies the input file (BACs have to be classified using the alphanumerical system, with the *Sal*I boxes of each repeat in brackets and each repeat separated by a tab), -o the output file, -t the temporary file, -a the type of analysis (0 for *Sal*I boxes and 1 for SNPs/InDels), -r for the randomization (0 = no randomization, 1 = randomization). The normalized association values were obtained by dividing the number of each rDNA association by the sum of the number of repeats taken into consideration. A list of all BACs with each rDNA unit categorized by their features is described in Suppl. Table S9.

### Monte Carlo simulation

The Monte Carlo simulation was performed running the Neighbour finder with an initial adaptation to shuffle the rDNA units within and between the BACs for 1000 times. The frequency of each rDNA association was calculated by summing the number of occurrences per association divided by 1000. We considered only frequencies larger than 0.90. The normalized association values were obtained by dividing the frequency of each rDNA association by the sum of the number of repeats taken into consideration.

#### Statistical Analysis

All statistical analyses, t Tests, Mann-Whitney test, and Pearson Correlation tests (as indicated in the figure legends) were performed using GraphPad Prism 8 software (GraphPad Software).

Unpaired, two-tailed Mann-Whitney tests were performed, since D’Agostino Pearson omnibus K2 normality testing revealed that most data were not sampled from a Gaussian population, and nonparametric tests were therefore required. Error bars indicate standard deviations.

## Data Availability

Accession numbers for all BACs are listed in #####. All sequencing data is deposited on the Sequencing Read Archive (SRA) under BioProject number #####. The whole genome sequencing data set from Kurzbauer et al. 2020 is deposited under BioProject number ####.

---

**Suppl. Fig. 1:**
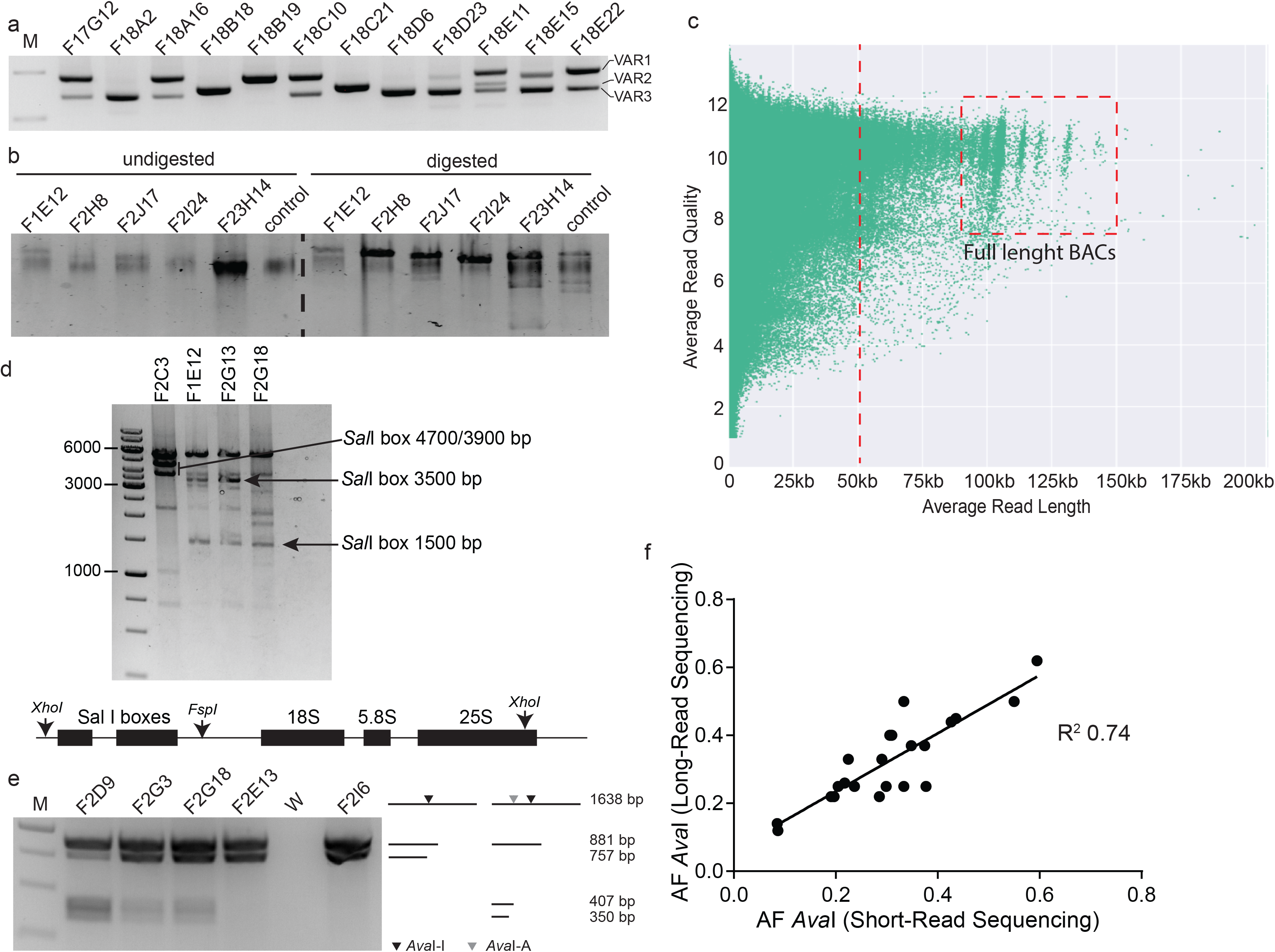
Sequencing and polishing BACs containing rDNA genes. **a**, Example of an agarose gel depicting the different 3’ETS variants (VAR 1-3) located within BACs. **b**, Example of a pulse field agarose gel with samples of BACs (“control” depicts BAC T15P10 that contains only one rDNA unit) before and after digestion with *Apa*I which linearizes the BACs without compromising the rDNA units. **c**, Plot showing an example of a NanoPore multiplex run. The individual reads are plotted according to average read quality vs. read length. Full length sequenced BACs can be visualized as clustered reads of ≥ 90 kb (red dashed box). Red dashed line marks the 50 kb threshold used for the assembly. **d**, Example of a restriction digest gel which confirms the correct assembly of the BACs F2C3, F1E12, F2G13, F2G18. The BACs were digested with *Xho*I and *Fsp*I that cut at the borders of the *Sal*I boxes and within the 25S rDNA. Arrows show the released *Sal*I boxes at the expected sizes. **e**, Example of an agarose gel showing the analysis of the presence/absence of the polymorphism at position 4133 (of the reference rDNA) that creates a CAPS marker that can be cleaved with *Ava*I. BACs F2D9, F2G3 and F2G18 contain the *Ava*I site, BACs F2E13 and F2I6 not. Black arrow heads indicate an *Ava*I site present in all rDNA units (*Ava*I-I), grey arrow heads indicate the cut site of the allelic *Ava*I site (*Ava*I-A). **f**, Correlation analysis of the *Ava*I allele frequencies derived from either long- or short-read sequencing approaches. Each dot represents an individual assembly.

**Suppl. Figure 2:**
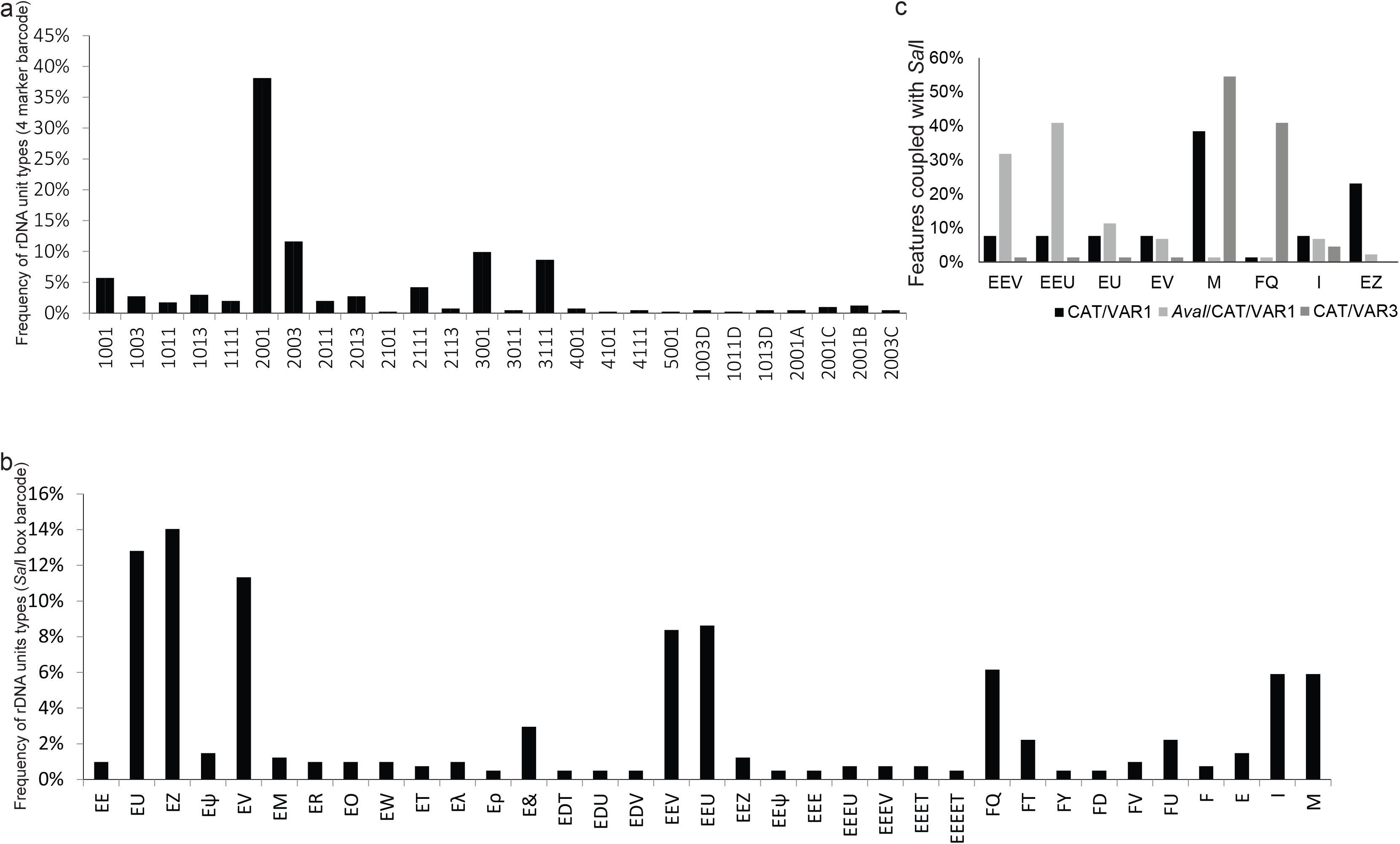
NOR2 contains a heterogeneous population of rDNA units. **a**, Graph depicting the percentage of rDNA units with the feature combinations: number of promoters (1 to 5), presence (1) or absence (0) of the *Ava*I restriction site, presence (1) or absence (0) of the CAT polymorphism and length of the 3’ ETS variant (1 or 3), presence of the deletion A-D. **b**, Percentage of rDNA variants carrying a certain *Sal*I box combination (see Table 1 for glossary). **c**, Percentage of rDNA variants 011 (black, presence of CAT and VAR1), 111 (dark gray, presence of *Ava*I, presence of CAT and VAR1) and 013 (light gray, presence of CAT and VAR3) associated with different *Sal*I box combinations (see Table 1 for glossary).

**Suppl. Figure 3:**
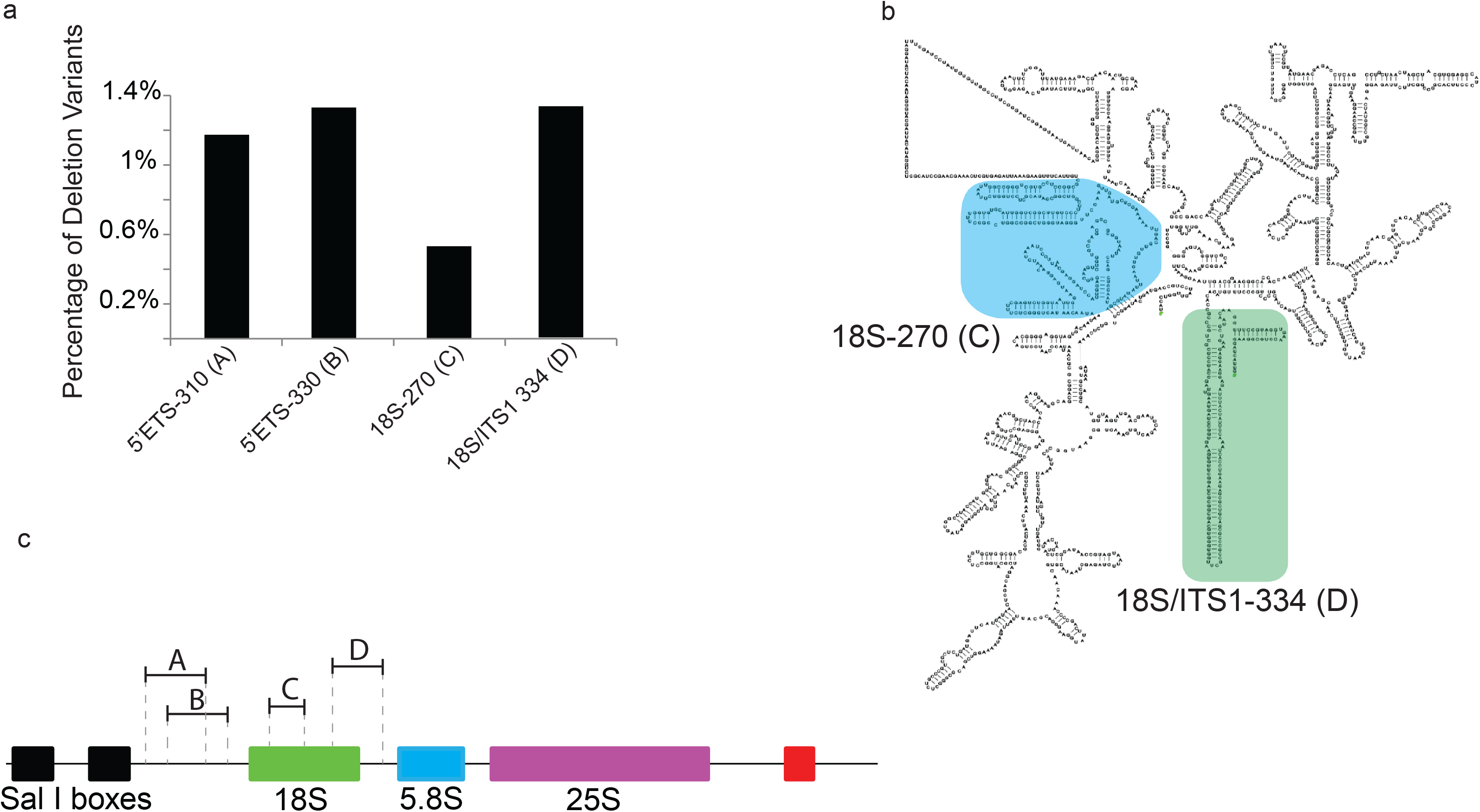
NOR2 contains a heterogeneous population of rDNA units. **a**, Percentage of rDNA deletion variants within the 5’ETS, 18S and spanning the 18S-ITS1 region. **b**, 2D structure of the 18S rRNA ^36^. Regions that would be theoretically lost in processed 18S rRNA transcribed from units encoding deletions are highlighted. Light blue shows the 270 bp deletion within the 18S ^18^ (“C”), light green shows the 334bp deletion that spans the 18S and the ITS1 (“D”). **c**, Graphical depiction of an rDNA unit with identified deletions indicated.

**Suppl. Figure 4:**
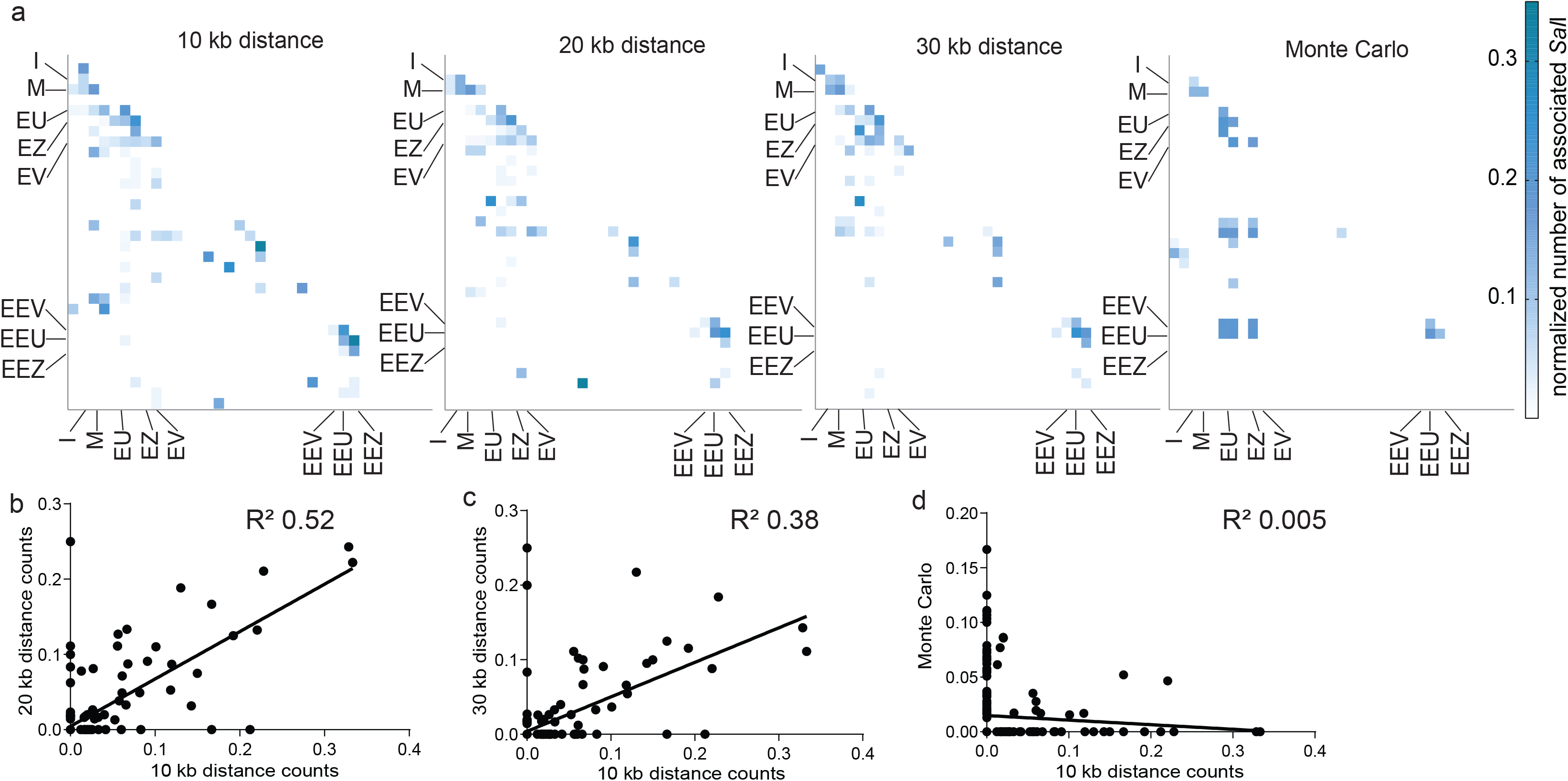
NOR2 is organized in distinct rDNA sub-clusters. **a**, Heat maps depicting the number of associated rDNA units carrying different *Sal*I box combinations at a 10kb (immediate neighbor), 20kb (neighbor one unit downstream) or 30kb (neighbor two units downstream) distance. The random control (10kb) was performed following a Monte Carlo simulation. Counts were normalized to the numbers of involved rDNA units. Blue color depicts the highest count (0.35) and whites the lowest (0). *Sal*I boxes are listed as follows for each heat map on the y-axis from top to bottom and for the x-axis from left to right: EE, EU, EZ, Eψ, EV, EM, ER, EO, EW, ET, Eλ, Eρ, E&, EDT, EDU, EDV, EEV, EEU, EEZ, EEψ, EEE, EEEU, EEEV, EEET, EEEET, FQ, FT, FY, FD, FV, FU, F, E, I, M (see Table S4 for glossary). **b-d**, Scatter plots showing the correlation between the frequencies of associated repeats carrying different *Sal*I box combinations at a 10 kb distance compared to: **b**, 20kb distance (R^2^=0.52); **c**, 30 kb distance (R^2^=0.38); **d,** Monte Carlo random control (R^2^=0.005).

## Table Legends

**Table 1: Glossary for the alphanumeric feature classification**

Glossary of the different features used to individualize each rDNA unit.

**Suppl. Table 1: Sequencing and polishing BACs containing rDNA contigs**

List of all BACs sequenced with their rDNA variant type, sample number, read cut-off size, number of reads used for the assembly and allelic frequency of two SNP/InDels (*Ava*I and CAT) derived from the Illumina sequencing (Illumina) and the long-read assembly (NanoPore)

**Suppl. Table 2: NOR2 contains a heterogeneous population of rDNA units**

List of all allelic frequencies per BAC occurring at positions relative to the reference rDNA repeat. Position (POS).

**Suppl. Table 3: Unequal distribution of SNP/InDels between NOR2 and NOR4**

List of the allelic frequencies, based on Illumina sequencing, occurring in the whole genome sequencing data set (Plant) and on NOR2 (BAC derived NOR2).

**Suppl. Table 4: Unequal distribution of SNP/InDels between NOR2 and NOR4**

List of all NOR2 called SNP/InDels with their position (POS), relative to the reference rDNA, the reference allelic variant (REF), the alternative variant (VAR), the quality threshold determined by LoFreq (QUAL) and the allelic frequency (AF)

**Suppl. Table 5: rRNA variants are differentially expressed between tissues**

List of the allelic frequencies, based on the ribosome capture RNA-seq and online retrieved polysome sequencing profiles, occurring along the 25S rRNA for each tissue analyzed. Positions (POS), whole genome (WG), adult leaves (AL), young leaves (YL), inflorescences (INFLO), siliques (S), adult leaves online data base (AL-DB), young leaves online data base (YL-DB), inflorescences online data base (INFLO-DB).

**Suppl. Table 6: rRNA variants are differentially expressed between tissues**

List of the allelic frequencies, based on the total RNA-seq and online retrieved total RNA-seq, occurring along the rDNA reference for each tissue analyzed. Positions (POS), whole genome (WG), adult leaves (AL), young leaves (YL), inflorescences (INFLO), siliques (S), adult leaves online data base (AL-DB), young leaves online data base (YL-DB), inflorescences online data base (INFLO-DB).

**Suppl. Table 7: BACs assemblies and accession number**

List of all BACs and their accession number as deposited on the NCBI server.

**Suppl. Table 8: Nanopore Runs**

List of all Nanopore sequencing runs and their details

**Suppl. Table 9: rDNA units categorized by their features**

List of all BACs with the units organized according to their position and categorized by the four feature combinations including the *Sal*I repeats.

**Suppl. File 1: rDNA units form sub-clusters**

Raw data used to generate the co-occurrence tables in Figure 2

**Suppl. File 2: rDNA reference repeat sequence**

Sequence of the reference rDNA used in this study

## Acknowledgment

We thank all members of the Schlögelhofer and Von Haeseler Labs for discussions. We are indebted to Luis Paulin for helping with hardware and software set-up. We thank Philip Ratzenböck for help with BAC DNA preparation and Melitta Münster for help with the ribosome pull-down. We are grateful to the ABRC stock centre for all BACs. We thank the Austrian Science Fund (PS: SFB F34, I 3685-B25, I1468, AI02955, AI03685; AvH: W1207-B09) for funding.

